# Genome engineering allows selective conversions of terephthalaldehyde to multiple valorized products in bacterial cells

**DOI:** 10.1101/2023.05.02.539072

**Authors:** Roman M. Dickey, Michaela A. Jones, Neil D. Butler, Ishika Govil, Aditya M. Kunjapur

## Abstract

Deconstruction of polyethylene terephthalate (PET) plastic waste generates opportunities for valorization to alternative products. We recently designed an enzymatic cascade that could produce terephthalaldehyde (TPAL) from terephthalic acid. Here, we showed that the addition of TPAL to growing cultures of *Escherichia coli* wild-type strain MG1655 and an engineered strain for reduced aromatic <u>a</u>ldehyde <u>re</u>ction (RARE) strain resulted in substantial reduction. We then investigated if we could mitigate this reduction using multiplex automatable genome engineering (MAGE) to create an *E. coli* strain with 10 additional knockouts in RARE. Encouragingly, we found this newly engineered strain enabled a 2.5-fold higher retention of TPAL over RARE after 24h. We applied this new strain for the production of *para*-xylylenediamine (pXYL) and observed a 6.8-fold increase in pXYL titer compared to RARE. Overall, our study demonstrates the potential of TPAL as a versatile intermediate in microbial biosynthesis of chemicals that derived from waste PET.

## Introduction

The increasing societal dependence on plastics derived from petroleum and natural gas feedstocks has generated a demand to divert plastic waste from landfills to alternative products^1–3^. As such, tremendous efforts have been made to enhance the sustainability and renewability in polymer life cycles. Particularly, polyethylene terephthalate (PET) plastic, one of the most commonly produced polyesters, has garnered increasing attraction with the development of chemical deconstruction through synthetic approaches^4–7^ or use of microbial PET-degrading enzymes^8–11^. Both enzymatic and chemical means of PET deconstruction toward the monomer unit terephthalic acid (TPA) generate opportunities to create higher value products or monomers for use in other classes of materials. In recent years, there have been several examples of platform chemicals produced biocatalytically from TPA, including gallic acid, pyrogallol, catechol, muconic acid, vanillic acid and vanillin made by live cells^12,13^. We recently identified carboxylic acid reductase enzymes that can efficiently generate the versatile dialdehyde terephthalaldehyde (TPAL) from TPA^14^. Our first step was to report the use of these enzymes *in vitro*, in part because it is unclear if these steps would function well in live cells given the potential instability and reactivity of TPAL^15^. However, the use of live cells to convert TPAL to useful chemical building blocks could provide a more cost effective and efficient biosynthetic process compared to the use of purified enzymes^16–18^.

Aldehydes can readily undergo numerous biological transformations to generate a large variety of potential products. Here, we envision utilizing the dialdehyde functionality of TPAL to access potential new opportunities in biosynthetic pathways, including asymmetrically functionalized products. Aldehyde-derived biosynthetic targets have ranged broadly^19^, including diamine polymer building blocks^14,20^, hydroxylated non-standard amino acids^21–25^, nitro alcohols^26^, and pharmaceutical mono-amine precursors^27–29^. Despite the biosynthetic possibilities available from aldehydes, their redox instability in cellular environments due to endogenous oxidoreductase activity remains a key challenge^30^. First, we were curious to learn if we could take advantage of the endogenous enzymes that reduce TPAL in growing wild-type *E. coli* K-12 MG1655 cells to form the target diol 1,4-benzenedimethanol (BDM), a valuable building block that can be utilized for the production of pesticides, perfumes, or dyes^31–34^. While this instability can be utilized to rapidly reduce aldehydes into alcohols products like BDM, the strongly reductive natural cellular environment mitigates the use of aldehydes as a platform intermediate. To address this challenge, alcohol dehydrogenases (ADHs) or aldo-keto reductases (AKRs) in *E. coli* can be deleted and have resulted in sustainable improvements in stability for a wide set of aromatic and aliphatic aldehydes under aerobic conditions^35–37^. Of particular interest, the RARE.Δ6 strain (more commonly known as the RARE strain), an *E. coli* MG1655 strain named for its reduced aromatic <u>a</u>ldehyde <u>re</u>duction, has alleviated the issue of aldehyde stability for many aromatic aldehydes^37^. Thus, we were also curious to learn if the RARE.Δ6 strain would allow us to convert TPAL to amines.

In this study, we investigated the stability of TPAL when supplemented to *E. coli* MG1655 and the RARE.Δ6 strain for the potential reduction by endogenous enzymes to the corresponding mono-or di-alcohols. We showed that TPAL is reduced by growing cell cultures of the MG1655 strain and, to our surprise, TPAL is also reduced by growing cell cultures of the RARE.Δ6 strain. Interestingly, we discovered that we could use the RARE.Δ6 strain to stably accumulate the mono-aldehyde mono-alcohol 4-(hydroxymethyl) benzaldehyde (4HMB), which can be used in polymer applications as a precursor that is converted to an aryl bromide on route to polymers bearing an aldehyde at their chain ends^38^. To determine if we could overcome reduction of either aldehyde functional group by identifying and inactivating additional aldehyde reductases, we used multiplex automatable genome engineering (MAGE) to perform translational gene inactivation of up to 10 additional ADH and AKR genes, partly guided by RNA-seq that revealed a previously unreported target whose expression was elevated by TPAL challenge. After performing these additional gene inactivations, we created a strain (RARE.Δ16) that achieved significant retention of TPAL under aerobic growth conditions after 24 h. Finally, we showed that the use of this strain can lead to large improvements in the biosynthesis of the diamine pXYL from TPAL. Our study thus exploits genome engineering and heterologous expression to demonstrate selective routes to three distinct building blocks from TPAL.

## Results

### TPAL is reduced in MG1655 and RARE.Δ6 under aerobic conditions

We first sought to measure the stability of TPAL and the potential to produce a diol product under aerobic growth conditions **(Fig. 1A)**. To evaluate whether reduction occurred during fermentation, we supplemented 5 mM TPAL to cultures of the *E. coli* MG1655 strain in LB media at mid-exponential phase. We grew cells in deep 96-well plates with 300 μL of media and used non-breathable aluminum seals to limit loss due to aldehyde volatility. We sampled the culture broth at two time points (4 h and 24 h after aldehyde addition) to gain some insight on the kinetics of aldehyde stability. In cultures of the wild-type *E. coli* MG1655 strain, we observed complete reduction of TPAL to BDM within 4 h, with this condition persisting after 24 h **(Fig. 1B)**. We expect that this reduction is catalyzed by several endogenous aldehyde reductases in *E. coli* that in prior studies we have worked to identify and eliminate for the stabilization of aldehydes^35–37,39^.

**Figure 1.**
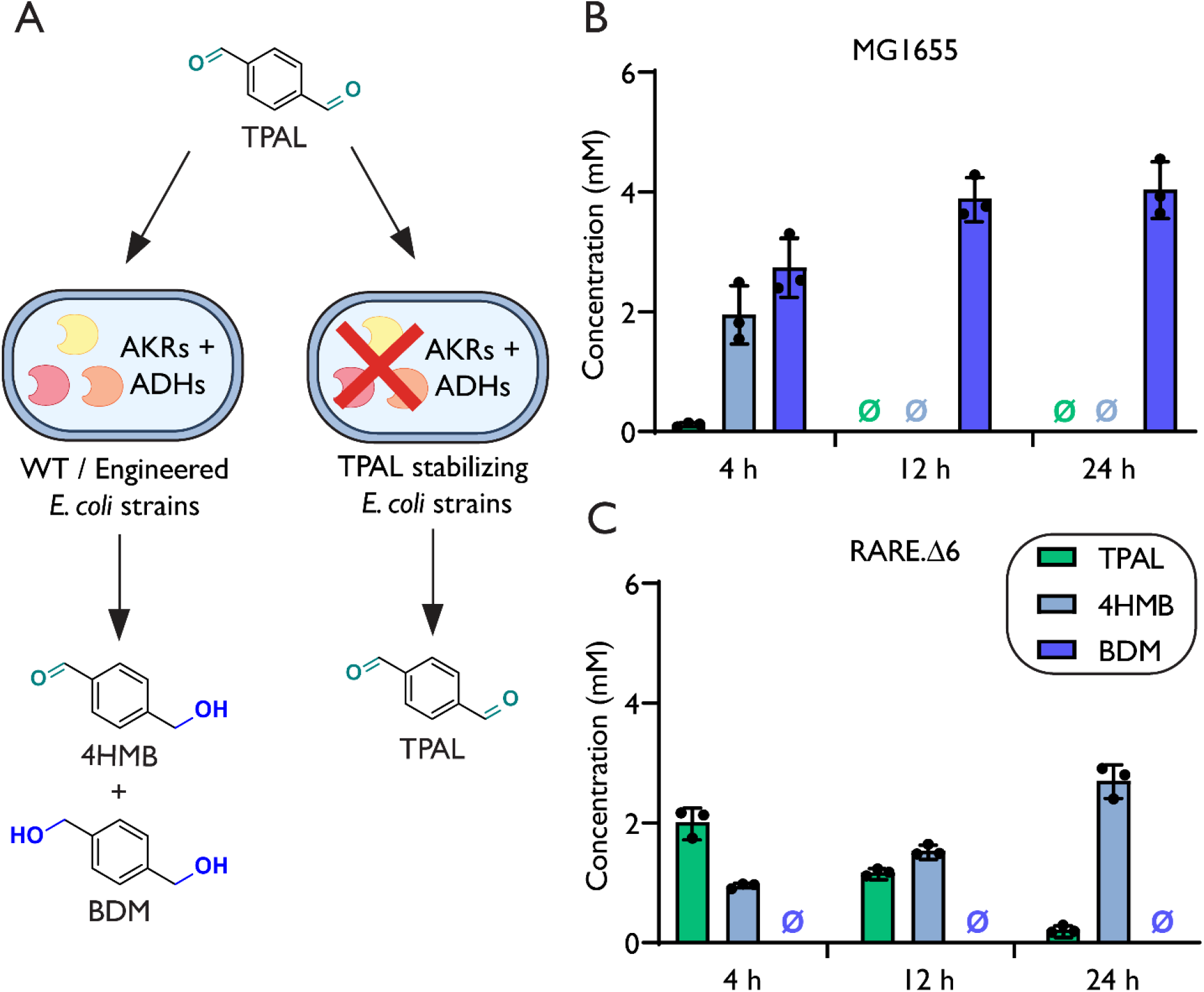
Evaluation of the stability of TPAL when supplemented to aerobic cultures of *E. coli*. **(A)** TPAL was added to culture media to determine its fate in the presence of growing *E. coli* cells over time. **(B)** Cultures of wild-type *E. coli* MG1655 were grown in LB media at 37°C and supplemented with 5 mM TPAL at mid-exponential phase. TPAL metabolites were tracked via HPLC at 4 h, 12 h and 24 h. **(C)** Cultures of the previously reported *E. coli* RARE.Δ6 strain were grown under identical conditions and TPAL metabolites were tracked via HPLC at 4 h, 12 h and 24 h. Data represents technical triplicates (n=3) where error bars represent the standard deviation across triplicates. Null sign indicates no detectable quantities were observed.

We were also curious about whether we could improve or enable the conversion of TPAL to other products by inactivating endogenous ADHs and AKRs. As a first step towards testing whether inactivation of aldehyde reductases could eliminate TPAL reduction, we evaluated TPAL stability in the previously engineered RARE.Δ6 strain under aerobic growth. Although we recently verified that the RARE.Δ6 strain reliably stabilizes a broad range of aromatic aldehydes under these conditions^40^, we were surprised to observe TPAL reduction at our first time point of 4 h with greater reduction seen at 24 h **(Fig. 1C)**. However, we observed that cultures of the RARE.Δ6 strain at 4 h and 24 h were able to stabilize the mono-aldehyde 4HMB, eliminating the complete reduction of TPAL to BDM. Thus, we can select production of BDM and 4HMB through the use of MG1655 and RARE.Δ6 respectively. While the RARE.Δ6 strain was able to provide enhanced TPAL stability over that of the wild-type MG1655 strain at 4 h, there is still significant reduction within our system at 24 h (0.22 ± 0.07 mM TPAL, 2.70 ± 0.27 mM 4HMB).

### Rational targeted gene inactivation enables TPAL stability

Given the effectiveness of combinatorial gene deletions at limiting the reduction of aldehydes, we next investigated whether we could mitigate the reduction of TPAL observed using additional genome engineering. We used MAGE to inactivate potential genes responsible for the reduction of TPAL **(Fig. 2A)**. Given the partial success of the RARE.Δ6 strain at increasing TPAL stability, we sought to identify additional ADH or AKR targets that could contribute to TPAL reduction. We first looked at additional ADH candidates reported in the previously engineered *E. coli* strain AL1728 which reported 13 aldehyde reductase deletions, several of which were not included in the RARE.Δ6 strain^36^. We generated an initial subset of 5 targeted gene deletions (S1: Δ*adhP* Δ*fucO* Δ*eutG* Δ*yiaY* Δ*adhE*) to test if additional ADH knockouts could provide enhanced TPAL stabilization. In addition to investigating aldehyde reducing enzymes that had been deleted in previous studies, we also sought to determine whether other, nonobvious targets existed. To do so, we performed an RNA-seq experiment comparing conditions with and without addition of TPAL. We grew RARE.Δ6 in culture tubes in M9-glucose media, where media composition was defined.

**Figure 2.**
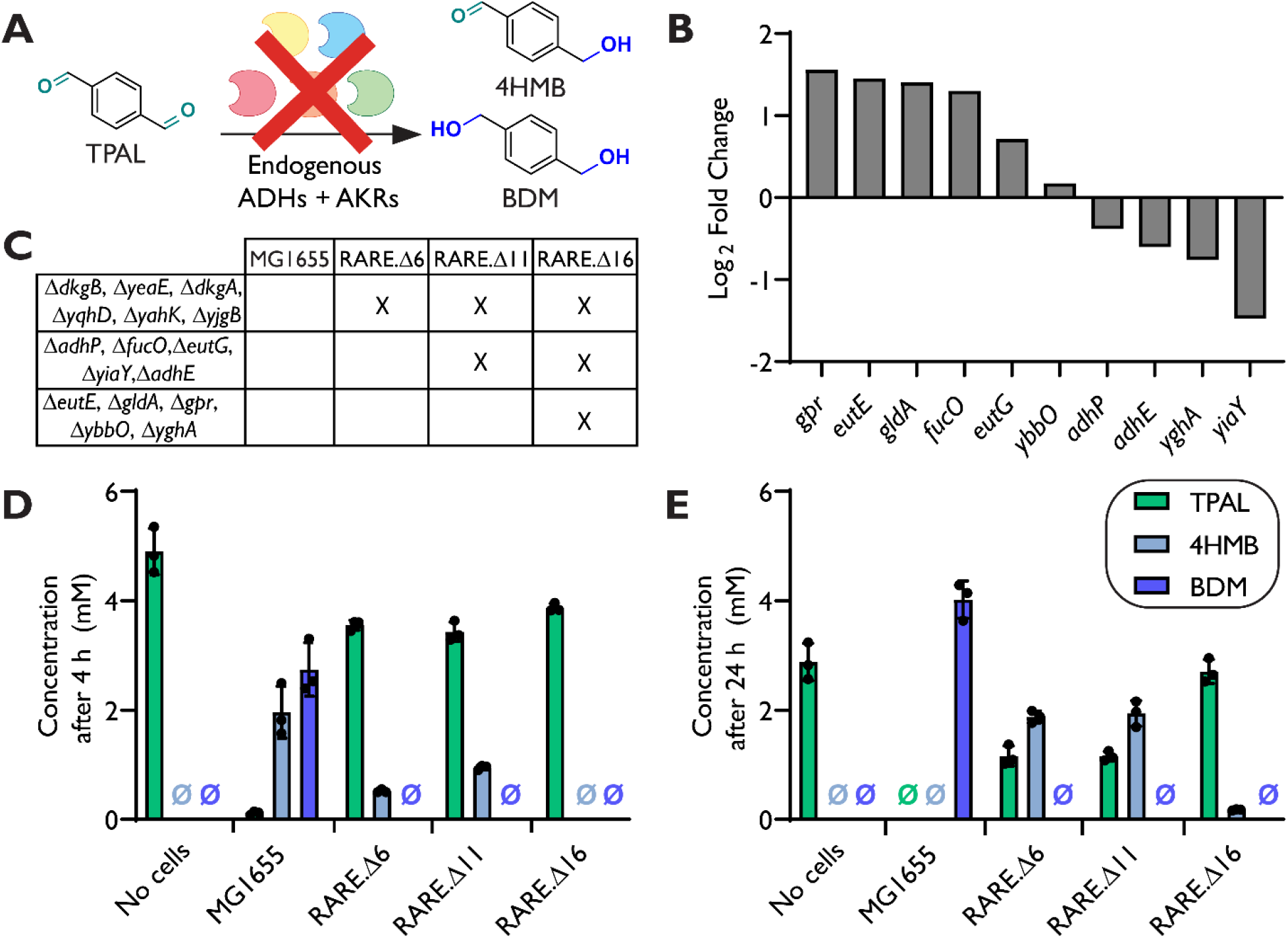
Genomic knockout of aldehyde reductases (ALR) toward improved TPAL stability in aerobic conditions. **(A)** Using the *E. coli* RARE.Δ6 strain as a basis, MAGE was performed to translationally knockout ALRs to generate two strains RARE.Δ11 (RARE.Δ6, Δ*adhP* Δ*fucO* Δ*eutG* Δ*yiaY* Δ*adhE*) and RARE.Δ16 (RARE.Δ11, Δ*eutE* Δ*gldA*, Δ*gpr* Δ*ybbO* Δ*yghA*). **(B)** Log_2_ fold change of targeted genes were determined through a TPAL challenge RNA-seq. Data represents TPAL supplementation in RARE.Δ6 compared to a RARE.Δ6 baseline in technical duplicates (n=2). **(C)** Gene knockouts contained within each strain. **(D&E)** Cultures of wild-type *E. coli* MG1655, RARE.Δ6, RARE.Δ11 and RARE.Δ16 were grown in LB media at 37°C and supplemented with 5 mM TPAL at mid-exponential phase. TPAL metabolites were tracked via HPLC at 4 h **(D)** and 24 h **(E)**. Data represents technical triplicates (n=3) where error bars represent the standard deviation across triplicates. Null sign indicates no detectable quantities were observed.

After cultures reached mid-exponential phase, we then either added 1 mM of TPAL and sealed culture tubes or simply sealed culture tubes and grew cultures for an additional 1.5 h. We then harvested and submitted total RNA for sequencing at Novogene. We then examined the sequencing results for upregulated ADH and AKR transcripts in the TPAL addition case. We selected *gpr, eutE* and *gldA* as targeted genes for deletion as all had relatively high fold changes (> 1) (**Fig 2B**). We also selected *ybbO* as we observed a small positive fold change. Additionally, we selected *yghA*, for deletion despite the down regulation as we still observed relatively high transcript levels. With this, we then constructed a second subset of 5 additional ADHs and AKRs for deletions (S2: Δ*eutE* Δ*gldA* Δ*gpr* Δ*ybbO* Δ*yghA*).

We utilized MAGE to inactivate S1 and S2 for a total of 10 gene deletions. We used 10 total rounds of MAGE to inactivate each subset of targeted genes in the *E. coli* RARE.Δ6 strain via introduction of in-frame stop codons within the first 100 codons. We then used multiplex allele-specific colony-PCR and confirmation by Sanger sequencing to obtain a variant containing the first subset of knockouts denoted as RARE.Δ11 (RARE.Δ6, S1) and a variant with all 10 knockouts denoted RARE.Δ16 (RARE.Δ11, S2) **(Fig 2C)**. Next, we sought to determine the stability of 5 mM TPAL under aerobic growth conditions in both new strains along with their progenitor strains. We were excited to observe no detectable reduction of TPAL after 4 h in RARE.Δ16 **(Fig. 2D)**. When compared to RARE.Δ6 at 24 h, we observed a 2.5-fold increase in TPAL concentration and a near 10-fold decrease in the 4HMB concentration **(Fig. 2E)**.

### Contribution of ALR knockouts to TPAL reduction

We set out to determine the impact of each of the additional 10 KOs on TPAL stability because of the potential tradeoffs on fitness and heterologous expression associated with multiple gene knockouts. To reintroduce each gene, we transformed RARE.Δ16 with a plasmid of each gene cloned from the wild type MG1655. We then cultured RARE.Δ6, RARE.Δ16, and distinct RARE.Δ16 transformants that overexpress each gene that had been targeted for inactivation under aerobic conditions at 37 ºC. At mid-exponential phase, we induced each culture and supplied 5 mM TPAL. To our surprise, here we observed that when each targeted gene was individually overexpressed, only 4 (*ybbO, gpr, yiay, yghA*) out of the 10 resulted in TPAL reduction **(Fig. 3A)**. Because MAGE is a rapid and combinatorial genome engineering strategy, we then chose to create a new strain to test the effect of introducing translational knockouts of only these four additional genes in RARE.Δ6, denoted as RARE.Δ10 (RARE.Δ6, Δ*ybbO* Δ*gpr* Δ*yiay* Δ*yghA*) **(Fig. 3B)**. We then evaluated the stability of TPAL in aerobic conditions with RARE.Δ10 and observed comparable stability to that of RARE.Δ16 **(Fig. 3C)**. This result indicated that the inactivation of only *ybbO, gpr, yiaY* and *yghA* were required to achieve the increased TPAL stability.

**Figure 3.**
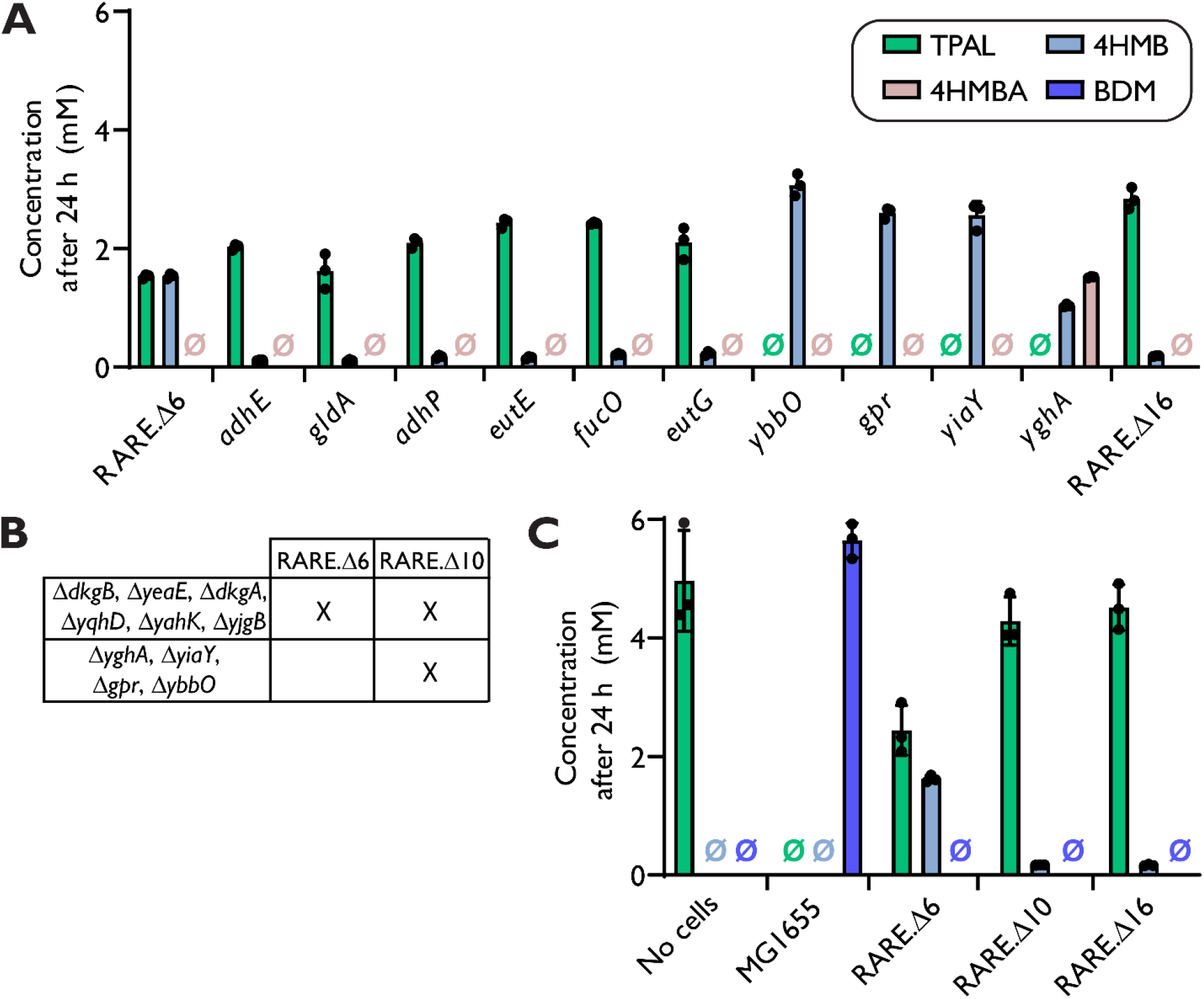
Evaluation of overexpressed ALR activity on TPAL stability. **(A)** Cultures of RARE.Δ6, RARE.Δ10 and RARE.Δ16 transformed with a single plasmid for each individual ALR gene from this study were grown in LB media at 37°C and supplemented with 5 mM TPAL and then induced of targeted gene at mid-exponential phase and tracked via HPLC after 24 h (4HMBA indicates 4-hydroxymethyl benzoic acid). **(B)** Gene knockouts contained within RARE.Δ10 strain. Using the *E. coli* RARE.Δ6 strain as a basis, MAGE was performed to translationally knockout the 4 ALRs that showed reduction of TPAL when overexpressed to generate the RARE.Δ10 strain (RARE.Δ6, Δ*ybbO* Δ*gpr* Δ*yiaY* Δ*yghA*). **(C)** Cultures of wild-type *E. coli* MG1655, RARE.Δ6, RARE.Δ10 and RARE.Δ16 were grown in LB media at 37°C and supplemented with 5 mM TPAL at mid-exponential phase. TPAL metabolites were tracked via HPLC at 24 h. Data represents technical triplicates (n=3) where error bars represent the standard deviation across triplicates. Null sign indicates no detectable quantities were observed.

### Characterization of the RARE.Δ10 and RARE.Δ16 strains for biocatalysis applications

With TPAL stability achieved, we then investigated whether the genome engineering performed impacted important parameters for functional application of these strains, namely effects on the cellular growth rate and protein overproduction. Thus, we investigated the growth rates across MG1655, RARE.Δ6, RARE.Δ10 and RARE.Δ16. We observed in both LB, a complex media, as well as MOPS EZ Rich-glucose, a defined media, there was no significant differences in the doubling time between the progenitor MG1655 strain and the engineered strains **(Fig. 4A, B, Table 1)**. However, we noticed there were small differences in growth rates towards late exponential and stationary phase, which resulted in lower final biomass concentrations of RARE.Δ16 for both media conditions. Furthermore, we used expression of a superfolder green fluorescent protein (sfGFP) reporter to compare protein production capacity of each previously mentioned strain. Each strain was transformed with a plasmid that harbored sfGFP and was grown in LB and MOPS EZ Rich-glucose media. We observed similar production of sfGFP when normalized by OD_600_ during exponential growth **(Fig. 4 C, D)**. Additionally, we noted that RARE.Δ16 had slightly increased normalized sfGFP in both media conditions. With the comparable levels in RARE.Δ10 and RARE.Δ16 to that of MG1655, our engineered strains show limited deleterious impact to growth rate and ability to overexpress desired protein despite the inactivation of up to 16 total genes.

**Table 1.** Growth performance of engineered TPAL retaining stains. Data represents average of technical triplicates (n=3) where error represent the standard deviation across triplicates.

|  | LB MEDIA |  | MOPS MEDIA |  |
| --- | --- | --- | --- | --- |
|  | Doubling time (min) | Final OD<sub>600</sub> | Doubling time (min) | Final OD<sub>600</sub> |
| MG1655 | 29.6 ± 1.5 | 0.66 ± 0.01 | 29.6 ± 1.5 | 0.66 ± 0.01 |
| RARE.Δ6 | 31.0 ± 0.4 | 0.64 ± 0.03 | 31.0 ± 0.4 | 0.64 ± 0.03 |
| RARE.Δ10 | 34.1 ± 0.7 | 0.71 ± 0.02 | 34.1 ± 0.7 | 0.71 ± 0.02 |
| RARE.Δ16 | 33.1 ± 0.5 | 0.55 ± 0.04 | 33.1 ± 0.5 | 0.55 ± 0.04 |

**Figure 4.**
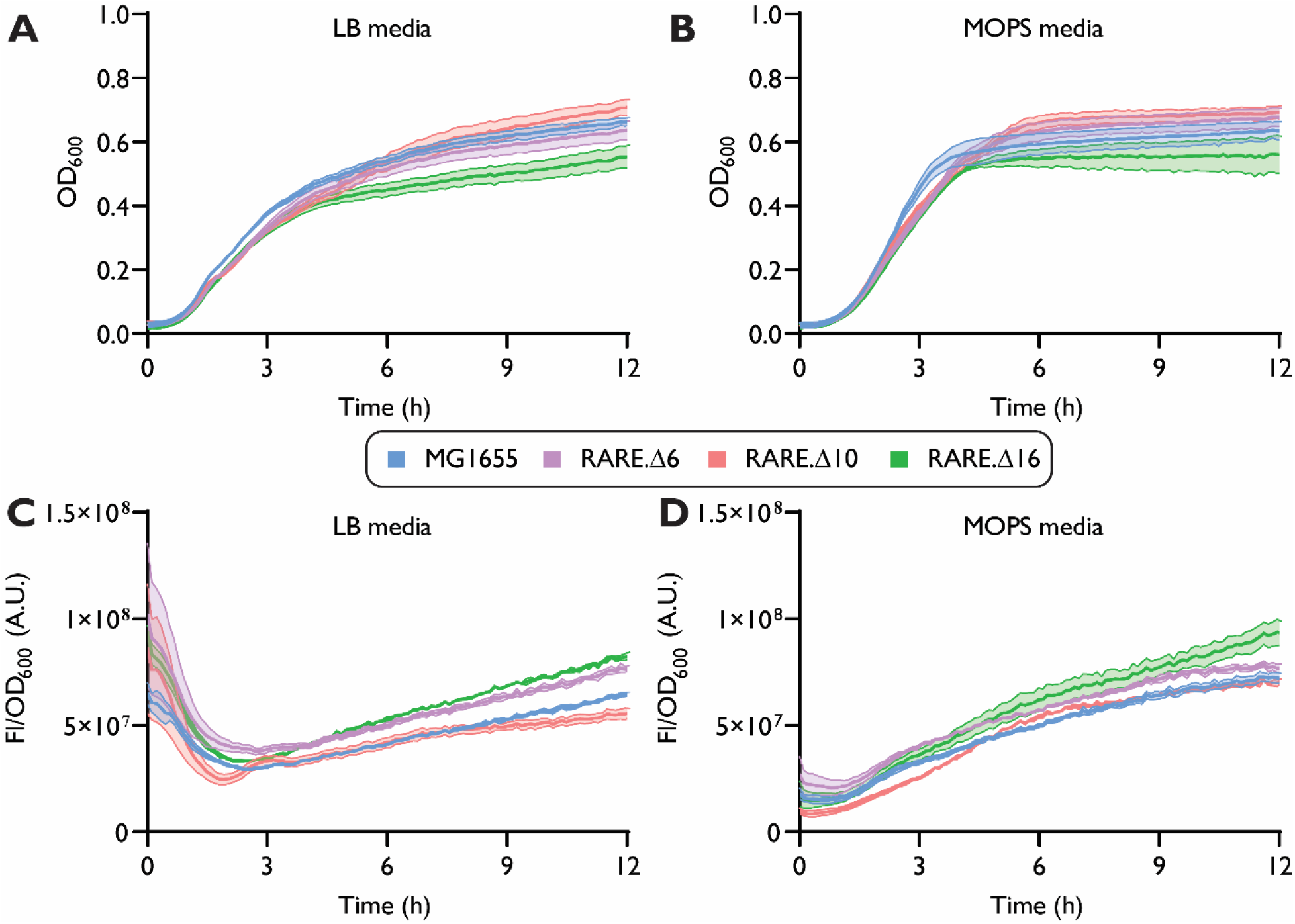
Growth and protein production performance of engineered TPAL retaining strains. Growth was monitored via optical density at 600 nm (OD_600_) in 96-well plate for 12 h in LB media **(A)** and in MOPS EZ Rich media **(B)**. Plasmid-based protein overexpression of superfolder green fluorescent protein (sfGFP) was monitored via 96-well plate in a plate reader for 12 h by measuring fluorescence (ex: 488 nm, em: 525 nm) normalized by OD_600_ in both LB media **(C)** and MOPS EZ Rich media **(D)** (A.U. indicates arbitrary units). Data represents technical triplicates (n=3) where error bars represent the standard deviation across triplicates.

### Application of the RARE.Δ16 strain for improved amine synthesis

We anticipate that the enhanced stability of TPAL in RARE.Δ16 affords opportunities to design potential biosynthesis pathways to convert TPAL to products besides BDM under aerobic fermentation conditions **(Fig. 5A)**. To do so, we looked into the enzyme class of ω-transaminases (TAs), which are reversible pyridoxal-5′-phosphate (PLP)-dependent enzymes that catalyze the transfer of an amino group between donor and acceptor^41^. Here, we evaluated the capability of RARE.Δ16 strain for improved biosynthesis of the diamine *para*-xylylenediamine (pXYL). pXYL can be utilized as a component in a wide variety of materials including polyamides, polyimides, or non-isocyanate polyurethanes^42,43^. We transformed MG1655, RARE.Δ6, and RARE.Δ16 with a plasmid construct that inducibly expresses a His6x-tagged TA from *Chromobacterium violaceum* (CvTA) and an L-alanine dehydrogenase for amino donor recycling from *Bacillus subtilis* (BsAlaDH). We cultured these strains under aerobic conditions in LB media with 400 μM PLP, 60 mM ammonium chloride and 100 mM L-alanine (amine donor). At mid-exponential phase (OD_600_ = 0.5-0.8), we induced each culture and supplied 5 mM TPAL. In MG1655, we observed the unexpected reduction of TPAL to 4HMB with no detectable reduction to BDM or amination to pXYL after 24 h **(Fig.5B)**. We hypothesize that the presence of the BsAlaDH could alter the co-factor pool and potentially limit further reduction towards BDM. We detected a similar result in RARE.Δ6 with the majority of TPAL reduced to 4HMB, however a small amount of pXYL was produced. We observed that the RARE.Δ16 strain achieved nearly 7-fold enhancement in pXYL production compared to the RARE.Δ6 stain after 24 h (1.93 ± 0.36 mM RARE.Δ16, 0.29 ± 0.20 mM RARE.Δ6). Thus, we have shown that the genetic engineering approach that led to the creation of RARE.Δ16 is able to unlock avenues to convert TPAL to multiple sets of valuable building block chemicals, including pXYL.

**Figure 5.**
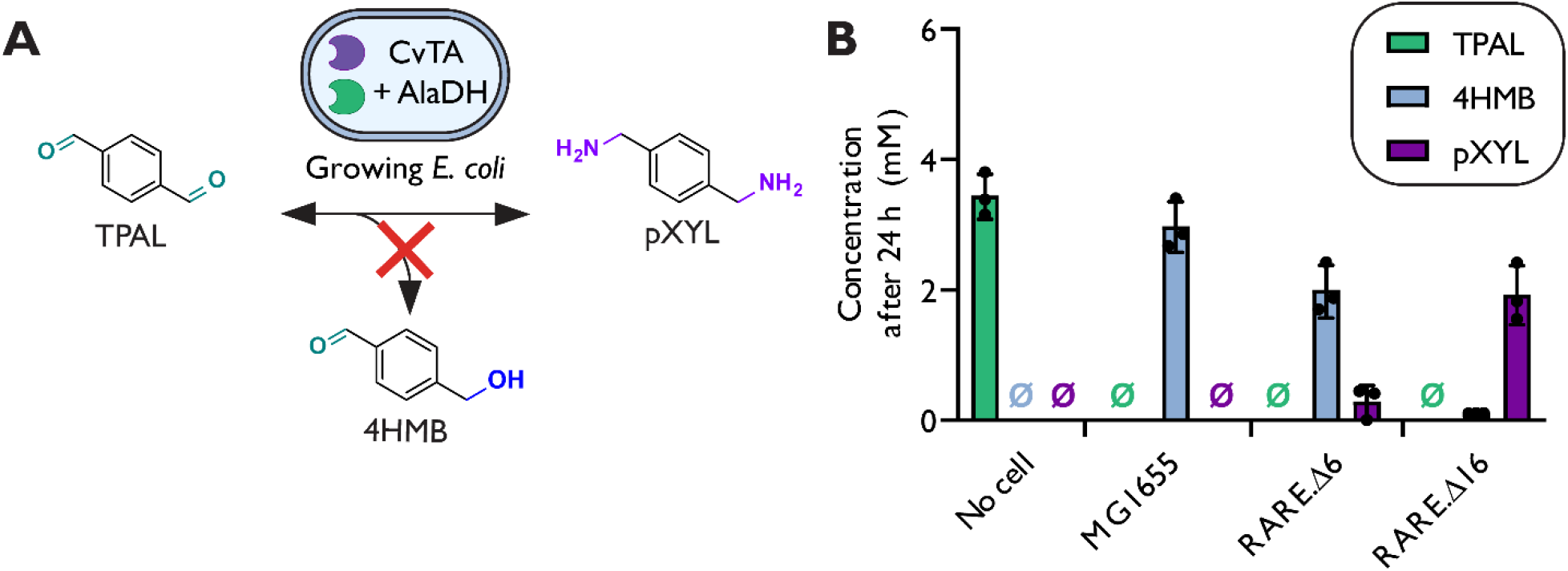
Biosynthesis of pXYL in engineered strains expressing CvTA. **(A)** We created *E. coli* strains containing CvTA and AlaDH that can convert TPAL to pXYL without additional reduction of TPAL. **(B)** Cultures of MG1655, RARE.Δ6 and RARE.Δ16 expressing CvTA were grown in LB media at 37°C and were supplemented with 100 mM L-alanine (amino donor) and 5 mM of TPAL at mid-exponential phase. TPAL, 4HMB, and pXYL concentrations were tracked via HPLC at 24 h. Data represents technical triplicates (n=3) where error bars represent the standard deviation across triplicates. Null sign indicates no detectable quantities were observed.

## Discussion

In this work, we identified that the wild-type MG1655 *E. coli* strain rapidly reduces TPAL to BDM in metabolically active cells. We then evaluated TPAL stability in the engineered RARE.Δ6 strain. Here, we showed that the RARE.Δ6 strain provided a 17.6-fold change in TPAL stability over that of the wild-type MG1655 after 4 h. However, at the later time point of 24 h, we observed the RARE.Δ6 strain had less than 8 ± 2% retention of TPAL. The RARE.Δ6 strain contains knockouts of six different genes that encode aldehyde reductases from two distinct families of enzymes AKR family and the NADH-dependent ADH family^37^. To determine if we could realize additional improvements in TPAL stability, we utilized the combinatorial genome engineering of MAGE to inactivate the translation of additional full-length ADHs and AKRs in the RARE.Δ6 strain. To identify potential genes that may be active on TPAL, we performed an RNAseq comparing conditions with and without supplementation of TPAL. With these genes identified and others identified in prior literature, we utilized MAGE to inactivate 10 additional genes into the RARE.Δ6 strain and deemed this strain RARE.Δ16 (*adhP, fucO, eutG, yiaY, adhE, eutE, gldA, gpr, ybbO, yghA*). We observed that the RARE.Δ16 strain had increased TPAL stability over MG1655 and RARE.Δ6 at short (4 h) and even at long time scales (24 h). We found that RARE.Δ16 contained 6 knockouts that were not required for TPAL stability and further showed that the inactivation of only *ybbO, gpr, yiay* and *yghA* combined with the RARE.Δ6 knockouts are necessary to achieve the TPAL stability exhibited in RARE.Δ16.

By performing differential expression analysis after an aldehyde challenge experiment, our study is the first to reveal the relevance of *gpr* to the reduction of aldehydes featured in engineered biosynthetic pathways. The *gpr* gene encoded the only AKR from our targeted knockouts. It has been shown to have high activity on methylglyoxal and has been utilized for glyoxal and methylglyoxal detoxification in *E. coli*^44–46^. Interestingly, *gpr* has been shown to have slight activity towards benzaldehyde and relatively high activity towards 4-nitrobenzaldehyde^44,45^. The *yiaY, yghA* and *ybbO* genes have been knocked out for a variety of short and long chain aliphatic reductions^35,36,47–50^. However, the activity of *yiaY* and *yghA* on aromatic aldehydes has not been shown, with *ybbO* very recently shown to contribute to the reduction of the aromatic aldehyde cinnamaldehyde^39^. Given what previous literature suggests, our work highlights the ability of RNAseq to reveal potential ADH and AKR that otherwise would not have been clear targets for gene inactivation for TPAL stability.

The improved stability of TPAL using live microbial cells can provide a more cost effective and efficient biosynthesis pathway over purified enzymes to other useful chemical building blocks^16–18^. A fermentative process circumvents the need for cell lysis and enzyme purification that can lower biocatalyst cost compared to purified enzymes^41,51–53^. This difference in biocatalyst cost can increase with multi-enzyme cascades and the need for additional co-factor regenerating enzymes. However, fermentative processes are limited by aldehyde toxicity to microbial cells. While aldehyde products could inhibit cellular growth, aldehyde intermediates could be kept at a low steady-state concentration with downstream enzymes.

Here, we demonstrated that the RARE.Δ16 strain can offer a significant improvement for TPAL biocatalysis using cells expressing CvTA. We observed that the RARE.Δ16 strain outperformed the RARE.Δ6 strain by nearly 7-fold after 24 h. There are several synthetic pathways that involve potential biosynthetic pathways involving TPAL as a substrate, intermediate, or product. In our previous work, we have shown potential biosynthesis pathways from PET deconstruction products like terephthalic acid or even mono-(2-hydroxyethyl) terephthalic acid to TPAL with the use of purified carboxylic acid reductases and a lipase. It is possible that other PLP-dependent enzymes could also utilize TPAL as a potential substrate. Our future work will look to identify whether RARE.Δ16 can serve as a platform strain to take advantage of the reactivity of aldehydes for the production of building blocks of diverse macromolecules, including non-standard amino acids^54^.

## Acknowledgements

We acknowledge support from the following funding source: The Center for Plastics Innovation, an Energy Frontier Research Center funded by the U.S. Department of Energy (DOE), Office of Science, Basic Energy Sciences, under Award No. # DE-SC0021166.

## Author Contributions

R.M.D designed and performed the MAGE to create RARE.Δ11. and RARE.Δ16, designed and conducted all stability experiments, analyzed data, prepared figures, and wrote the manuscript; M.A.J. initially documented ADH activity on TPAL, designed and performed MAGE to create RARE.Δ11. N.D.B performed the TPAL challenge and RNA-seq experiment as well as genome sequencing of RARE.Δ16. I.G. helped designed and performed MAGE to create RARE.Δ11. A.M.K. contributed to writing the manuscript, secured funding, provided oversight, and reviewed the manuscript.

## Conflict of Interest Statement

None

Additional information including Materials and Methods can be found in the Supporting Information document available online.

## Notes

### Competing Interest Statement

The authors have declared no competing interest.

