## Supplemental Information for "Genome engineering allows selective conversions of terephthalaldehyde to multiple valorized products in bacterial cells"

### I. Supplementary Tables

**Table S1.** Strains and plasmids used in this study.

| Name | Relevant genotype | Source |
|---|---|---|
| | ***E. coli* strains** | |
| DH5α | F– Φ80*lac*ZΔM15 Δ(*lac*ZYA-argF) U169 *rec*A1 *end*A1 *hsd*R17 (rK–, mK+) *pho*A *sup*E44 λ– *thi*-1 *gyr*A96 *rel*A1 | NEB |
| MG1655 | F- λ- *ilv*G- *rfb-50 rph-1* | ATCC 700926 |
| MG1655 (DE3) | F- λ- *ilv*G- *rfb-50 rph-1* (λ DE3)<br>λ DE3 = λ sBamHIo ΔEcoRI-B int::(*lac*I::PlacUV5::T7 gene1) i21 Δ*nin*5 | Previous study[1] |
| RARE.Δ6 | MG1655(DE3) Δ*dkgB* Δ*yeaE* Δ*(yqhC-dkgA)* Δ*yahK* Δ*yjgB* | Previous study [1] |
| RARE.Δ6-MAGE | RARE.Δ6 harboring pORTMAGE-Ec1 | This study |
| RARE.Δ11 | RARE.Δ6 Δ*adhP,* Δ*fucO,* Δ*eutG,* Δ*yiaY,* Δ*adhE* | This study |
| RARE.Δ16 | RARE.Δ6 Δ*adhP,* Δ*fucO,* Δ*eutG,* Δ*yiaY,* Δ*adhE,* Δ*eutE,* Δ*gldA,* Δ*gpr,* Δ*ybbO,* Δ*yghA* | This study |
| RARE.Δ10 | RARE.Δ6 Δ*yiaY,* Δ*gpr,* Δ*ybbO,* Δ*yghA* | This study |
| RMD001 | MG1655(DE3) harboring pZE-Ub-sfGFP | This study |
| RMD002 | RARE.Δ6 harboring pZE-Ub-sfGFP | This study |
| RMD003 | RARE.Δ10 harboring pZE-Ub-sfGFP | This study |
| RMD004 | RARE.Δ16 harboring pZE-Ub-sfGFP | This study |
| RMD005 | RARE.Δ16 harboring pACYC-adhP | This study |
| RMD006 | RARE.Δ16 harboring pACYC-fucO | This study |
| RMD007 | RARE.Δ16 harboring pACYC-eutG | This study |
| RMD008 | RARE.Δ16 harboring pACYC-yiaY, | This study |
| RMD009 | RARE.Δ16 harboring pACYC-adhE | This study |
| RMD010 | RARE.Δ16 harboring pACYC-eutE | This study |
| RMD011 | RARE.Δ16 harboring pACYC-gldA | This study |
| RMD012 | RARE.Δ16 harboring pACYC-gpr | This study |
| RMD013 | RARE.Δ16 harboring pACYC-ybbO | This study |
| RMD014 | RARE.Δ16 harboring pACYC-yghA | This study |
| RMD015 | RARE.Δ16 harboring pACYC-CvTA-AlaDH | This study |
| | **Plasmids** | |
| pORTMAGE-EC1 | RSF1010 ori, $Kan^R$, harboring CspRecT and *mutL* (E32K) genes | Previous study [2] |
| pZE-Ub-sfGFP | ColE1 ori, $Kan^R$, TetR, Tet promoter with a codon optimized ubiquitin fused to a codon-optimized superfolder GFP | This study |
| pACYC-adhP | p15a ori, $Cm^R$, LacI, Lac promoter with the gene for *E. coli* ADH *adhP* subcloned off of the genome. | This study |
| pACYC-fucO | p15a ori, $Cm^R$, LacI, Lac promoter with the gene for *E. coli* ADH *fucO* subcloned off of the genome. | This study |
| pACYC-eutG | p15a ori, $Cm^R$, LacI, Lac promoter with the gene for *E. coli* ADH *eutG* subcloned off of the genome. | This study |
| pACYC-yiaY, | p15a ori, $Cm^R$, LacI, Lac promoter with the gene for *E. coli* ADH *yiaY,* subcloned off of the genome. | This study |
| pACYC-adhE | p15a ori, $Cm^R$, LacI, Lac promoter with the gene for *E. coli* ADH *adhE* subcloned off of the genome. | This study |
| pACYC-eutE | p15a ori, $Cm^R$, LacI, Lac promoter with the gene for *E. coli* ADH *eutE* subcloned off of the genome. | This study |
| pACYC-gldA | p15a ori, $Cm^R$, LacI, Lac promoter with the gene for *E. coli* ADH *gldA* subcloned off of the genome. | This study |
| pACYC-gpr | p15a ori, $Cm^R$, LacI, Lac promoter with the gene for *E. coli* AKR *gpr* subcloned off of the genome. | This study |
| pACYC-ybbO | p15a ori, $Cm^R$, LacI, Lac promoter with the gene for *E. coli* ADH *ybbO* subcloned off of the genome. | This study |
| pACYC-yghA | p15a ori, $Cm^R$, LacI, Lac promoter with the gene for *E. coli* ADH *yghA* subcloned off of the genome. | This study |
| pACYC-CvTA-AlaDH | P15a ori, $Cm^R$, LacI, Lac promoter with the codon-optimized gene for ω-transaminase from *Chromobacterium violaceum* and a separate Lac promoter with the codon-optimized gene for alanine dehydrogenase from *Bacillus subtilis* | This study |

**Table S2.** Oligonucleotides used in this study (bold and underline nucleotide represent MAGE inserted mutation).

| Oligo Name | Sequence (5' to 3') |
|---|---|

| | |
|---|---|
| pacyc-adhE-fwd | CTTTAATAAGGAGATATACCATGGCTGTTACTAATGTCGCTG |
| pacyc-adhe-rev | GATCCCCCATCAAGCTTTTAAGCGGATTTTTTCGCTTTTTTC |
| pacyc-adhp-fwd | CTTTAATAAGGAGATATACCATGAAGGCTGCAGTTGTTACG |
| pacyc-adhp-rev | GATCCCCCATCAAGCTTTTAGTGACGGAAATCAATCACCATG |
| pacyc-bb-fwd-rareko | AAGCTTGATGGGGGATC |
| pacyc-bb-rev-rareko | GGTATATCTCCTTATTAAAGTTAAACAA |
| pacyc-eute-fwd | CTTTAATAAGGAGATATACCATGAATCAACAGGATATTGAACAGG |
| pacyc-eutE-rev | GATCCCCCATCAAGCTTTTAAACAATGCGAAACGCATC |
| pacyc-eutg-fwd | CTTTAATAAGGAGATATACCATGCAAAATGAATTGCAGACCG |
| pacyc-eutg-rev | GATCCCCCATCAAGCTTTTATTGCGCCGCTGCG |
| pacyc-fuco-fwd | CTTTAATAAGGAGATATACCATGGCTAACAGAATGATTCTGAAC |
| pacyc-fuco-rev | GATCCCCCATCAAGCTTTTACCAGGCGGTATGGTAAAG |
| pacyc-gldA-fwd | CTTTAATAAGGAGATATACCATGGACCGCATTATTCAATCAC |
| pacyc-glda-rev | GATCCCCCATCAAGCTTTTATTCCCACTCTTGCAGGAAAC |
| pacyc-gpr-fwd | CTTTAATAAGGAGATATACCATGGTCTGGTTAGCGAATCC |
| pacyc-gpr-rev | GATCCCCCATCAAGCTTTCATTTATCGGAAGACGCCTG |
| pacyc-ybbo-fwd | CTTTAATAAGGAGATATACCATGACTCATAAAGCAACGGAGATC |
| pacyc-ybbo-rev | GATCCCCCATCAAGCTTTCACCCCTGCAATATTTTGTCC |
| pacyc-yghA-fwd | CTTTAATAAGGAGATATACCATGTCTCATTTAAAAGACCCGACC |
| pacyc-ygha-rev | GATCCCCCATCAAGCTTTTAACCTAAATGCTCGCCGC |
| pacyc-yiay-fwd | CTTTAATAAGGAGATATACCATGGCAGCTTCAACGTTCTTTATTC |
| pacyc-yiay-rev | GATCCCCCATCAAGCTTTTACATCGCTGCGCGATAAATC |
| pacyc-cvta-ins-fwd | ACAGCCAGGATCCGAATTCGATGCAGAAGCAGCGTAC |
| pacyc-cvta-ins-rev | GGGGATCCCCCATCAAGCTTTCAGGCAAGTCCGCGAG |
| pacyc_bb_fwd | AGCTTGATGGGGGATCC |
| pacyc_bb_rev | CGAATTCGGATCCTGGC |
| pacyc-bb-cvta-ala-rev | ATGTATATCTCCTTCTTATACTTAACTAATATACTAAG |
| pacyc-bb-cvta-ala-fwd | TTAACCTAGGCTGCTGCC |
| pacyc-cvta-aladh-ala-fwd | TATAAGAAGGAGATATACATATGGGCAGCAGCCATC |
| pacyc-cvta-aladh-ala-rev | CAGCAGCCTAGGTTAATTACGCGCCCGCCA |
| Gpr-ko-oligo | TCGCTCGGTTTATGGCACAATTTCGGTCACGTTAACGCGCTG**T**AAT**G**A**T**AGCGTGCGATCCTGCGTAAAGC GTTTGATTTGGGCATTACG |
| EutE-ko-oligo | ATGGCAGCAATGGCTAACTGGCGCATTGCCACGCTTTTTAAtC**A**TT**A**CT**A**GGCGACTTTGGCTGCCGCAACG GCGTCATCCAGGGACGCG |
| YbbO-ko-oligo | TGACTCATAAAGCAACGGAGATCCTGACAGGTAAAGTTATG**T**AA**TG**AT**A**GGTCTTAATTACCGGATGTTCC AGTGGAATTGGCCTGGAAA |
| GldA-ko-oligo | AAGATGCTGGACTGGTAGTAGAAATTGCGCCGTTTGGCGGT**T**AATG**A**T**A**GCAAAATGAGATCGACCGTCTG CGTGGCATCGCGGAGACTG |
| YghA-ko-oligo | CCCTTAAAGGACGTTGAACCTGAAGGGAGAAAAACGATGTCT**T**A**G**T**G**A**T**AAGACCCGACCACGCAGTATT ACACTGGTGAATATCCCAAA |
| adhP-ko-oligo | AGCGTTTCGTTACCACTGTTACAGTATTCGCAATGACCGCATC**A**TT**A**GTAGAACCACGCCACGCTGGCACG ATCGCCTGGTTTTAATGAG |
| fucO-ko-oligo | TTTATCGGTGACGATCAGCGCCTTCTG**T**TA**T**C**A**ACGGCGTTTCACCTCATCGGTTAAAGCCCCAACAGCACC CCGACCAAACCATGCCGT |
| eutG-ko-oligo | CTGCCATCACGAACAGATGTTTCAGCCCACGCGTTTGCGCTT**A**CT**A**Tt**A**GCAACTGCTCACCGAGCCCGGAC CGCAAAGCGTCACCGGTG |
| yiaY-ko-oligo | GCTATCGCAATTATTCTCTTTAAGT**C**ATT**AAT**AACCTGCGGCGACGTTTTCCGTGGTGGGGTTAGGTTGGGT GCCATCATAAATAACGCT |
| adhE-ko-oligo | TCGCTGAACTTAACGCACTCGTAGAGCGTGTAAAAAAAGCCCAG**T**G**AT**AATATGCCAGTTTCACTCAAGAG CAAGTAGACAAAATCTTCC |
| Gpr-fwd-wt | TTTCGGTCACGTTAACGCGCTGG |
| Gpr-fwd-mut | CAATTTCGGTCACGTTAACGCGCTGT |
| EutE-fwd-wt | CTGGCGCATTGCCACGCTTTTTAAC |
| EutE-fwd-mut | ACTGGCGCATTGCCACGCTTTTTAAT |
| YbbO-fwd-wt | CAATTCCACTGGAACATCCGGTAATTAAGACCG |
| YbbO-fwd-mut | CCAATTCCACTGGAACATCCGGTAATTAAGACCT |
| GldA-fwd-wt | CGCAGACGGTCGATCTCATTTTGCG |
| GldA-fwd-mut | ACGCAGACGGTCGATCTCATTTTGCT |
| YghA-fwd-wt | CGTTGAACCTGAAGGGAGAAAAACGATGTCTC |

| | |
|---|---|
| YghA-fwd-mut | CGTTGAACCTGAAGGGAGAAAAACGATGTCTT |
| Gpr-rev-200 | CACATATCGTAGCCAGCCTTGGTAGAGATAATCAG |
| EutE-rev-300 | TGTTACTGGTGAGTGGCAGTTCCGC |
| YbbO-rev-400 | GCAATGACGGTTTGCGTGGTTTTCAGC |
| GldA-rev-500 | ACCGCGTAGATGCCCAGGGC |
| YghA-rev-600 | GACTTGGCTGGTATGCCTGGATTGACG |
| EutE-fwd-wt-retry | CGCATTGCCACGCTTTTTAAT |
| EutE-mut-fwd-retry | CGCATTGCCACGCTTTTTAAC |
| EutE-rev-retry | GTGAGTGGCAGTTCCGC |
| gpr-fwd-seq | GGTCTGGTTAGCGAATCCCGAACGTTAC |
| EutE-fwd-seq | CAAATTTATCTTCAACGCGCCCCATGCC |
| YbbO-fwd-seq | CCGGTAAATCCCATGCTGTTCATGCGC |
| GldA-fwd-seq | CCGGAACACCCATGAAATGTGCCAGTG |
| YghA-fwd-seq | AGCACGCAACAGCAAATCAACATATACGTTAAACC |
| eutG-400-rev | AAGCGCGGTCACGGTGAAG |
| eutG-fwd-mut | CCCACGCGTTTGCGCTTA |
| eutG-fwd-wt | CCACGCGTTTGCGCTTG |
| eutG-fwd-seq | ATGGCGATACCTTTAACGGTC |
| yiaY-300-rev | GAGTGTGTTATGGCAGCTTCA |
| YiaY-fwd-mut | CGCTATCGCAATTATTCTCTTTAAGTC |
| YiaY-fwd-wt | ACGCTATCGCAATTATTCTCTTTAAGTA |
| YiaY-fwd-seq | CGCTGATTCATTGACTGATG |
| adhE-fwd-wt | GATTTTGTCTACTTGCTCTTGAGTGAAACTGGCATATTC |
| adhE-fwd-mut | GATTTTGTCTACTTGCTCTTGAGTGAAACTGGCATATTAT |
| adhE-rev | ACAATTTATTAACTGTTAGCTATAATGGCGAAAAGCGATGC |
| adhE-seq-fwd | TACTCTCGTATTCGAGCAGATG |
| adhP-rev | GTGTTGTGGTGTATGTCATACCGATCT |
| adhP-mut-fwd | CAGTATTCGCAATGACCGCATCa |
| adhP-WT-fwd | CAGTATTCGCAATGACCGCATCc |
| adhP-seq-fwd | ACCGATCTTCATGTTAAGAATG |
| fucO-fwd-wt | GTGACGATCAGCGCCTTCTGA |
| fucO-fwd-mut | GTGACGATCAGCGCCTTCTGt |
| fucO-rev-150 | AGTAATTTCGTAAAGCAACAAGGAGAAGGATG |
| fucO-seq-fwd | GACGACAGTAATTGTTGGGTTG |

**Table S3.** Sequences of proteins expressed in this paper.

| Oligo Name | DNA CDS | Protein Sequence |
|---|---|---|
| pZE-Ub-GFP | ATGCAGATTTTTGTGAAGACTTTAACAGGTAAGACGATTACCCTGGAGGTGG<br>AGTCCTCGGACACCATCGATAATGTAAAATCAAAAATCCAAGATAAGGAAG<br>GAATCCCTCCAGACCAGCAACGTCTGATTTTCGCAGGTAAACAACTGGAGG<br>ATGGTCGCACGCTTTCGGACTACAACATCCAGAAAGAATCTACCCTTCATTT<br>GGTTCTGCGTCTGCGTGGAGGATAGTTGTTTGTGCAGGAGCTTGCATCCAAG<br>GGCGAGGAGCTCTTTACTGGCGTAGTACCAATTCTCGTAGAGCTCGATGGCG<br>ATGTAAATGGCCATAAGTTTTCCGTACGCGGCGAGGGCGAGGGCGATGCAA<br>CTAACGGCAAGCTCACTCTCAAGTTTATTTGTACTACTGGCAAGCTCCCAGT<br>ACCATGGCCAACTCTCGTAACTACTCTGACCTATGGCGTACAATGTTTTTCCC<br>GCTATCCAGATCACATGAAGCAACATGATTTTTTTAAGTCCGCAATGCCAGA<br>GGGCTATGTACAAGAGCGCACTATTAGCTTTAAGGATGATGGCACCTATAA<br>GACTCGCGCAGAGGTAAAGTTTGAGGGCGATACTCTCGTAAATCGCATTGA<br>GCTCAAGGGCATTGATTTTAAGGAGGATGGCAATATTCTCGGCCATAAGCTG<br>GAGTATAATTTCAATTCCCATAATGTATACATTACCGCAGATAAGCAAAAGA<br>ATGGCATTAAGGCGAATTTTAAGATTCGCCATAATGTGGAGGATGGCTCCGT<br>ACAACTCGCAGATCATTATCAACAAAATACTCCAATTGGCGATGGCCCAGTA<br>CTCCTCCCAGATAATCATTATCTCTCCACTCAATCCGTGCTCTCCAAAGATCC<br>AAATGAGAAGCGCGATCACATGGTACTCCTGGAGTTTGTAACTGCAGCAGG<br>CATTACTCATGGCATGGATGAGCTCTATAAGCTCGAGCACCACCACCACCAC<br>CACCACTAA | MQIFVKTLTGKTITLEVES<br>SDTIDNVKSKIQDKEGIPP<br>DQQRLIFAGKQLEDGRTL<br>SDYNIQKESTLHLVLRLR<br>GGYLFVQELASKGEELFT<br>GVVPILVELDGDVNGHKF<br>SVRGEGEGDATNGKLTLK<br>FICTTGKLPVPWPTLVTTL<br>TYGVQCFSRYPDHMKQH<br>DFFKSAMPEGYVQERTISF<br>KDDGTYKTRAEVKFEGD<br>TLVNRIELKGIDFKEDGNI<br>LGHKLEYNFNSHNVYITA<br>DKQKNGIKANFKIRHNVE<br>DGSVQLADHYQQNTPIGD<br>GPVLLPDNHYLSTQSVLS<br>KDPNEKRDHMVLLEFVT<br>AAGITHGMDELYKLEHH<br>HHHHH |
| pACYC-CvTA-AlaDH | CvTA:<br>ATGGGCAGCAGCCATCACCATCATCACCACAGCCAGGATCCGAATTCGATG<br>CAGAAGCAGCGTACAACATCGCAATGGCGCGAACTTGACGCCGCTCATCAC<br>CTGCATCCCTTCACCGATACCGCCTCCCTTAACCAGGCCGGCGCGCGCGTGA<br>TGACACGTGGAGAAGGGGTGTATTTGTGGGACTCGGAGGGAAATAAAATCA<br>TCGACGGTATGGCTGGATTATGGTGTGTGAACGTTGGCTACGGTCGTAAGGA | CvTA:<br><br>MGSSHHHHHHSQDPNSM<br>QKQRTTSQWRELDAAHH<br>LHPFTDTASLNQAGARVM |

| | | |
|---|---|---|
| | CTTTGCCGAAGCGGCCCGTCGTCAGATGGAAGAATTACCGTTCTACAATACT<br>TTTTTCAAAACAACCCATCCTGCGGTCGTAGAGTTATCTTCATTATTGGCGG<br>AAGTCACTCCAGCAGGGTTTGACCGCGTGTTTTATACAAATAGTGGATCAGA<br>ATCGGTTGACACAATGATCCGTATGGTCCGTCGTTACTGGGACGTCCAAGGC<br>AAACCGGAGAAGAAGACGTTAATCGGCCGCTGGAATGGTTATCACGGTTCG<br>ACCATTGGAGGTGCATCTCTTGGGGGCATGAAGTATATGCATGAGCAGGGT<br>GATTTGCCTATCCCTGGCATGGCGCACATCGAACAACCGTGGTGGTATAAGC<br>ACGGTAAAGACATGACGCCGGACGAGTTTGGAGTTGTCGCTGCGCGTTGGTT<br>GGAAGAGAAGATCCTGGAAATTGGGGCGGACAAGGTAGCCGCCTTCGTAGG<br>AGAACCAATCCAAGGTGCCGGGGGAGTGATCGTCCCGCCAGCTACCTATTG<br>GCCCGAGATCGAGCGCATTTGCCGTAAATATGACGTATTGCTGGTTGCAGAT<br>GAGGTAATTTGTGGCTTCGGGCGCACCGGGGAGTGGTTCGGGCACCAACAT<br>TTCGGTTTTCAGCCGGACTTATTTACGGCGGCGAAGGGTTTAAGCTCAGGTT<br>ATTTACCGATTGGGGCTGTGTTTGTGGGCAAGCGTGTTGCCGAAGGCTTAAT<br>CGCGGGAGGCGACTTTAATCACGGATTCACATACTCTGGACACCCGGTTTGT<br>GCCGCAGTAGCTCACGCGAATGTAGCCGCATTACGTGACGAGGGAATCGTC<br>CAGCGTGTGAAGGACGATATCGGCCCTTATATGCAGAAGCGCTGGCGCGAG<br>ACTTTTTCACGTTTTGAGCACGTAGACGATGTGCGTGGCGTAGGCATGGTAC<br>AGGCCTTTACCTTAGTCAAAAATAAAGCTAAGCGCGAGTTGTTCCCAGACTT<br>TGGCGAAATCGGAACGTTGTGTCGCGATATCTTTTTTCGCAATAATCTTATC<br>ATGCGCGCTTGCGGGGATCATATTGTAAGTGCCCCGCCATTGGTGATGACTC<br>GTGCCGAGGTAGATGAGATGTTAGCAGTCGCAGAGCGCTGCCTTGAGGAGT<br>TTGAGCAAACATTAAAAGCTCGCGGACTTGCCTGA<br>AlaDH:<br>ATGGGCAGCAGCCATCACCATCATCACCACATGATCATAGGGGTTCCTAAA<br>GAGATAAAAAACAATGAAAACCGTGTCGCATTAACACCCGGGGGCGTTTCT<br>CAGCTCATTTCAAACGGCCACCGGGTGCTGGTTGAAACAGGCGCGGGCCTT<br>GGAAGCGGATTTGAAAATGAAGCCTATGAGTCAGCAGGAGCGGAAATCATT<br>GCTGATCCGAAGCAGGTCTGGGACGCCGAAATGGTCATGAAAGTAAAAGAA<br>CCGCTGCCGGAAGAATATGTTTATTTTCGCAAAGGACTTGTGCTGTTTACGT<br>ACCTTCATTTAGCAGCTGAGCCTGAGCTTGCACAGGCCTTGAAGGATAAAGG<br>AGTAACTGCCATCGCATATGAAACGGTCAGTGAAGGCCGGACATTGCCTCTT<br>CTGACGCCAATGTCAGAGGTTGCGGGCAGAATGGCAGCGCAAATCGGCGCT<br>CAATTCTTAGAAAAGCCTAAAGGCGGAAAAGGCATTCTGCTTGCCGGGGTG<br>CCTGGCGTTTCCCGCGGAAAAGTAACAATTATCGGAGGAGGCGTTGTCGGG<br>ACAAACGCGGCGAAAATGGCTGTCGGCCTCGGTGCAGATGTGACGATCATT<br>GACTTAAACGCAGACCGCTTGCGCCAGCTTGATGACATCTTCGGCCATCAGA<br>TTAAAACGTTAATTTCTAATCCGGTCAATATTGCTGATGCTGTGGCGGAAGC<br>GGATCTCCTCATTTGCGCGGTATTAATTCCGGGTGCTAAAGCTCCGACTCTT<br>GTCACTGAGGAAATGGTAAAACAAATGAAACCCGGTTCAGTTATTGTTGAT<br>GTAGCGATCGACCAAGGCGGCATCGTCGAAACTGTCGACCATATCACAACA<br>CATGATCAGCCAACATATGAAAAACACGGGGTTGTGCATTATGCTGTAGCG<br>AACATGCCAGGCGCAGTCCCTCGTACATCAACAATCGCCCTGACTAACGTTA<br>CTGTTCCATACGCGCTGCAAATCGCGAACAAAGGGGCAGTAAAAGCGCTCG<br>CAGACAATACGGCACTGAGAGCGGGTTTAAACACCGCAAACGGACACGTGA<br>CCTATGAAGCTGTAGCAAGAGATCTAGGCTATGAGTATGTTCCTGCCGAGAA<br>AGCTTTACAGGATGAATCATCTGTGGCGGGTGCTTAA | TRGEGVYLWDSEGNKIID<br>GMAGLWCVNVGYGRKD<br>FAEAARRQMEELPFYNTF<br>FKTTHPAVVELSSLLAEVT<br>PAGFDRVFYTNSGSESVD<br>TMIRMVRRYWDVQGKPE<br>KKTLIGRWNGYHGSTIGG<br>ASLGGMKYMHEQGDLPIP<br>GMAHIEQPWWYKHGKD<br>MTPDEFGVVAARWLEEKI<br>LEIGADKVAAFVGEPIQG<br>AGGVIVPPATYWPEIERIC<br>RKYDVLLVADEVICGFGR<br>TGEWFGHQHFGFQPDLFT<br>AAKGLSSGYLPIGAVFVG<br>KRVAEGLIAGGDFNHGFT<br>YSGHPVCAAVAHANVAA<br>LRDEGIVQRVKDDIGPYM<br>QKRWRETFSRFEHVDDVR<br>GVGMVQAFTLVKNKAKR<br>ELFPDFGEIGTLCRDIFFRN<br>NLIMRACGDHIVSAPPLV<br>MTRAEVDEMLAVAERCL<br>EEFEQTLKARGLA*<br><br>AlaDH:<br>MGSSHHHHHHMIIGVPKE<br>IKNNENRVALTPGGVSQLI<br>SNGHRVLVETGAGLGSGF<br>ENEAYESAGAEIIADPKQV<br>WDAEMVMKVKEPLPEEY<br>VYFRKGLVLFTYLHLAAE<br>PELAQALKDKGVTAIAYE<br>TVSEGRTLPLLTPMSEVA<br>GRMAAQIGAQFLEKPKGG<br>KGILLAGVPGVSRGKVTII<br>GGGVVGTNAAKMAVGL<br>GADVTIIDLNADRLRQLD<br>DIFGHQIKTLISNPVNIADA<br>VAEADLLICAVLIPGAKAP<br>TLVTEEMVKQMKPGSVIV<br>DVAIDQGGIVETVDHITTH<br>DQPTYEKHGVVHYAVAN<br>MPGAVPRTSTIALTNVTV<br>PYALQIANKGAVKALAD<br>NTALRAGLNTANGHVTY<br>EAVARDLGYEYVPAEKA<br>LQDESSVAGA* |
| pACYC-adhe | ATGGCTGTTACTAATGTCGCTGAACTTAACGCACTCGTAGAGCGTGTAAAAA<br>AAGCCCAGCGTGAATATGCCAGTTTCACTCAAGAGCAAGTAGACAAAATCT<br>TCCGCGCCGCCGCTCTGGCTGCTGCAGATGCTCGAATCCCACTCGCGAAAAT<br>GGCCGTTGCCGAATCCGGCATGGGTATCGTCGAAGATAAAGTGATCAAAAA<br>CCACTTTGCTTCTGAATATATCTACAACGCCTATAAAGATGAAAAAACCTGT<br>GGTGTTCTGTCTGAAGACGACACTTTTGGTACCATCACTATCGCTGAACCAA<br>TCGGTATTATTTGCGGTATCGTTCCGACCACTAACCCGACTTCAACTGCTATC<br>TTCAAATCGCTGATCAGTCTGAAGACCCGTAACGCCATTATCTTCTCCCCGC<br>ACCCGCGTGCAAAAGATGCCACCAACAAAGCGGCTGATATCGTTCTGCAGG<br>CTGCTATCGCTGCCGGTGCTCCGAAAGATCTGATCGGCTGGATCGATCAACC<br>TTCTGTTGAACTGTCTAACGCACTGATGCACCACCCAGACATCAACCTGATC<br>CTCGCGACTGGTGGTCCGGGCATGGTTAAAGCCGCATACAGCTCCGGTAAA<br>CCAGCTATCGGTGTAGGCGCGGGCAACACTCCAGTTGTTATCGATGAAACTG<br>CTGATATCAAACGTGCAGTTGCATCTGTACTGATGTCCAAAACCTTCGACAA<br>CGGCGTAATCTGTGCTTCTGAACAGTCTGTTGTTGTTGTTGACTCTGTTTATG<br>ACGCTGTACGTGAACGTTTTGCAACCCACGGCGGCTATCTGTTGCAGGGTAA<br>AGAGCTGAAAGCTGTTCAGGATGTTATCCTGAAAAACGGTGCGCTGAACGC<br>GGCTATCGTTGGTCAGCCAGCCTATAAAATTGCTGAACTGGCAGGCTTCTCT<br>GTACCAGAAAACACCAAGATTCTGATCGGTGAAGTGACCGTTGTTGATGAA<br>AGCGAACCGTTCGCACATGAAAAACTGTCCCCGACTCTGGCAATGTACCGC | MAVTNVAELNALVERVK<br>KAQREYASFTQEQVDKIF<br>RAAALAAADARIPLAKM<br>AVAESGMGIVEDKVIKNH<br>FASEYIYNAYKDEKTCGV<br>LSEDDTFGTITIAEPIGIICG<br>IVPTTNPTSTAIFKSLISLK<br>TRNAIIFSPHPRAKDATNK<br>AADIVLQAAIAAGAPKDLI<br>GWIDQPSVELSNALMHHP<br>DINLILATGGPGMVKAAY<br>SSGKPAIGVGAGNTPVVID<br>ETADIKRAVASVLMSKTF<br>DNGVICASEQSVVVVDSV<br>YDAVRERFATHGGYLLQ<br>GKELKAVQDVILKNGALN<br>AAIVGQPAYKIAELAGFS<br>VPENTKILIGEVTVVDESE<br>PFAHEKLSPTLAMYRAKD<br>FEDAVEKAEKLVAMGGIG |

| | | |
|---|---|---|
| | GCTAAAGATTTCGAAGACGCGGTAGAAAAAGCAGAGAAACTGGTTGCTATG<br>GGCGGTATCGGTCATACCTCTTGCCTGTACACTGACCAGGATAACCAACCGG<br>CTCGCGTTTCTTACTTCGGTCAGAAAATGAAAACGGCGCGTATCCTGATTAA<br>CACCCCAGCGTCTCAGGGTGGTATCGGTGACCTGTATAACTTCAAACTCGCA<br>CCTTCCCTGACTCTGGGTTGTGGTTCTTGGGGTGGTAACTCCATCTCTGAAAA<br>CGTTGGTCCGAAACACCTGATCAACAAGAAAACCGTTGCTAAGCGAGCTGA<br>AAACATGTTGTGGCACAAACTTCCGAAATCTATCTACTTCCGCCGTGGCTCC<br>CTGCCAATCGCGCTGGATGAAGTGATTACTGATGGCCACAAACGTGCGCTCA<br>TCGTGACTGACCGCTTCCTGTTCAACAATGGTTATGCTGATCAGATCACTTCC<br>GTACTGAAAGCAGCAGGCGTTGAAACTGAAGTCTTCTTCGAAGTAGAAGCG<br>GACCCGACCCTGAGCATCGTTCGTAAAGGTGCAGAACTGGCAAACTCCTTCA<br>AACCAGACGTGATTATCGCGCTGGGTGGTGGTTCCCCGATGGACGCCGCGA<br>AGATCATGTGGGTTATGTACGAACATCCGGAAACTCACTTCGAAGAGCTGG<br>CGCTGCGCTTTATGGATATCCGTAAACGTATCTACAAGTTCCCGAAAATGGG<br>CGTGAAAGCGAAAATGATCGCTGTCACCACCACTTCTGGTACAGGTTCTGAA<br>GTCACTCCGTTTGCGGTTGTAACTGACGACGCTACTGGTCAGAAATATCCGC<br>TGGCAGACTATGCGCTGACTCCGGATATGGCGATTGTCGACGCCAACCTGGT<br>TATGGACATGCCGAAGTCCCTGTGTGCTTTCGGTGGTCTGGACGCAGTAACT<br>CACGCCATGGAAGCTTATGTTTCTGTACTGGCATCTGAGTTCTCTGATGGTC<br>AGGCTCTGCAGGCACTGAAACTGCTGAAAGAATATCTGCCAGCGTCCTACC<br>ACGAAGGGTCTAAAAATCCGGTAGCGCGTGAACGTGTTCACAGTGCAGCGA<br>CTATCGCGGGTATCGCGTTTGCGAACGCCTTCCTGGGTGTATGTCACTCAAT<br>GGCGCACAAACTGGGTTCCCAGTTCCATATTCCGCACGGTCTGGCAAACGCC<br>CTGCTGATTTGTAACGTTATTCGCTACAATGCGAACGACAACCCGACCAAGC<br>AGACTGCATTCAGCCAGTATGACCGTCCGCAGGCTCGCCGTCGTTATGCTGA<br>AATTGCCGACCACTTGGGTCTGAGCGCACCGGGCGACCGTACTGCTGCTAAG<br>ATCGAGAAACTGCTGGCATGGCTGGAAACGCTGAAAGCTGAACTGGGTATT<br>CCGAAATCTATCCGTGAAGCTGGCGTTCAGGAAGCAGACTTCCTGGCGAAC<br>GTGGATAAACTGTCTGAAGATGCATTCGATGACCAGTGCACCGGCGCTAAC<br>CCGCGTTACCCGCTGATCTCCGAGCTGAAACAGATTCTGCTGGATACCTACT<br>ACGGTCGTGATTATGTAGAAGGTGAAACTGCAGCGAAGAAAGAAGCTGCTC<br>CGGCTAAAGCTGAGAAAAAAGCGAAAAAATCCGCTTAA | HTSCLYTDQDNQPARVSY<br>FGQKMKTARILINTPASQ<br>GGIGDLYNFKLAPSLTLG<br>CGSWGGNSISENVGPKHL<br>INKKTVAKRAENMLWHK<br>LPKSIYFRRGSLPIALDEVI<br>TDGHKRALIVTDRFLFNN<br>GYADQITSVLKAAGVETE<br>VFFEVEADPTLSIVRKGAE<br>LANSFKPDVIIALGGGSPM<br>DAAKIMWVMYEHPETHF<br>EELALRFMDIRKRIYKFPK<br>MGVKAKMIAVTTTSGTGS<br>EVTPFAVVTDDATGQKYP<br>LADYALTPDMAIVDANLV<br>MDMPKSLCAFGGLDAVT<br>HAMEAYVSVLASEFSDGQ<br>ALQALKLLKEYLPASYHE<br>GSKNPVARERVHSAATIA<br>GIAFANAFLGVCHSMAHK<br>LGSQFHIPHGLANALLICN<br>VIRYNANDNPTKQTAFSQ<br>YDRPQARRRYAEIADHLG<br>LSAPGDRTAAKIEKLLAW<br>LETLKAELGIPKSIREAGV<br>QEADFLANVDKLSEDAFD<br>DQCTGANPRYPLISELKQI<br>LLDTYYGRDYVEGETAA<br>KKEAAPAKAEKKAKKSA<br>* |
| pACYC-adhP | ATGAAGGCTGCAGTTGTTACGAAGGATCATCATGTTGACGTTACGTATAAAA<br>CACTGCGCTCACTGAAACATGGCGAAGCCCTGCTGAAAATGGAGTGTTGTG<br>GTGTATGTCATACCGATCTTCATGTTAAGAATGGCGATTTTGGTGACAAAAC<br>CGGCGTAATTCTGGGCCATGAAGGCATCGGTGTGGTGGCAGAAGTGGGTCC<br>AGGTGTCACCTCATTAAAACCAGGCGATCGTGCCAGCGTGGCGTGGTTCTAC<br>GAAGGATGCGGTCATTGCGAATACTGTAACAGTGGTAACGAAACGCTCTGC<br>CGTTCAGTTAAAAATGCCGGATACAGCGTTGATGGCGGGATGGCGGAAGAG<br>TGCATCGTGGTCGCCGATTACGCGGTAAAAGTGCCAGATGGTCTGGACTCGG<br>CGGCGGCCAGCAGCATTACCTGTGCGGGAGTCACCACCTACAAAGCCGTTA<br>AGCTGTCAAAAATTCGTCCAGGGCAGTGGATTGCTATCTACGGTCTTGGCGG<br>TCTGGGTAACCTCGCCCTGCAATACGCGAAGAATGTCTTTAACGCCAAAGTG<br>ATCGCCATTGATGTCAATGATGAGCAGTTAAAACTGGCAACCGAAATGGGC<br>GCAGATTTAGCGATTAACTCACACACCGAAGACGCCGCCAAAATTGTGCAG<br>GAGAAAACTGGTGGCGCTCACGCTGCGGTGGTAACAGCGGTAGCTAAAGCT<br>GCGTTTAACTCGGCAGTTGATGCTGTCCGTGCAGGCGGTCGTGTTGTGGCTG<br>TCGGTCTACCGCCGGAGTCTATGAGCCTGGATATCCCACGTCTTGTGCTGGA<br>TGGTATTGAAGTGGTCGGTTCGCTGGTCGGCACGCGCCAGGATTTAACTGAA<br>GCCTTCCAGTTTGCCGCCGAAGGTAAAGTGGTGCCGAAAGTCGCCCTGCGTC<br>CGTTAGCGGACATCAACACCATCTTTACTGAGATGGAAGAAGGCAAAATCC<br>GTGGCCGCATGGTGATTGATTTCCGTCACTAA | MKAAVVTKDHHVDVTY<br>KTLRSLKHGEALLKMECC<br>GVCHTDLHVKNGDFGDK<br>TGVILGHEGIGVVAEVGP<br>GVTSLKPGDRASVAWFYE<br>GCGHCEYCNSGNETLCRS<br>VKNAGYSVDGGMAEECI<br>VVADYAVKVPDGLDSAA<br>ASSITCAGVTTYKAVKLS<br>KIRPGQWIAIYGLGGLGN<br>LALQYAKNVFNAKVIAID<br>VNDEQLKLATEMGADLAI<br>NSHTEDAAKIVQEKTGGA<br>HAAVVTAVAKAAFNSAV<br>DAVRAGGRVVAVGLPPES<br>MSLDIPRLVLDGIEVVGSL<br>VGTRQDLTEAFQFAAEGK<br>VVPKVALRPLADINTIFTE<br>MEEGKIRGRMVIDFRH* |
| pACYC-eutE | ATGAATCAACAGGATATTGAACAGGTGGTGAAAGCGGTACTGCTGAAAATG<br>CAAAGCAGTGACACGCCGTCCGCCGCCGTTCATGAGATGGGCGTTTTCGCGT<br>CCCTGGATGACGCCGTTGCGGCAGCCAAAGTCGCCCAGCAAGGGTTAAAAA<br>GCGTGGCAATGCGCCAGTTAGCCATTGCTGCCATTCGTGAAGCAGGCGAAA<br>AACACGCCAGAGATTTAGCGGAACTTGCCGTCAGTGAAACCGGCATGGGGC<br>GCGTTGAAGATAAATTTGCAAAAAACGTCGCTCAGGCGCGCGGCACACCAG<br>GCGTTGAGTGCCTCTCTCCGCAAGTGCTGACTGGCGACAACGGCCTGACCCT<br>AATTGAAAACGCACCCTGGGGCGTGGTGGCTTCGGTGACGCCTTCCACTAAC<br>CCGGCGGCAACCGTAATTAACAACGCCATCAGCCTGATTGCCGCGGGCAAC<br>AGCGTCATTTTTGCCCCGCATCCGGCGGCGAAAAAAGTCTCCCAGCGGGCG<br>ATTACGCTGCTCAACCAGGCGATTGTTGCCGCAGGTGGGCCGGAAAACTTAC<br>TGGTTACTGTGGCAAATCCGGATATCGAAACCGCGCAACGCTTGTTCAAGTT<br>TCCGGGTATCGGCCTGCTGGTGGTAACCGGCGGCGAAGCGGTAGTAGAAGC<br>GGCGCGTAAACACACCAATAAACGTCTGATTGCCGCAGGCGCTGGCAACCC<br>GCCGGTAGTGGTGGATGAAACCGCCGACCTCGCCCGTGCCGCTCAGTCCATC<br>GTCAAAGGCGCTTCTTTCGATAACAACATCATTTGTGCCGACGAAAAGGTAC<br>TGATTGTTGTTGATAGCGTAGCCGATGAACTGATGCGTCTGATGGAAGGCCA<br>GCACGCGGTGAAACTGACCGCAGAACAGGCGCAGCAGCTGCAACCGGTGTT | MNQQDIEQVVKAVLLKM<br>QSSDTPSAAVHEMGVFAS<br>LDDAVAAAKVAQQGLKS<br>VAMRQLAIAAIREAGEKH<br>ARDLAELAVSETGMGRV<br>EDKFAKNVAQARGTPGV<br>ECLSPQVLTGDNGLTLIEN<br>APWGVVASVTPSTNPAAT<br>VINNAISLIAAGNSVIFAPH<br>PAAKKVSQRAITLLNQAI<br>VAAGGPENLLVTVANPDI<br>ETAQRLFKFPGIGLLVVTG<br>GEAVVEAARKHTNKRLIA<br>AGAGNPPVVVDETADLA<br>RAAQSIVKGASFDNNIICA<br>DEKVLIVVDSVADELMRL<br>MEGQHAVKLTAEQAQQL<br>QPVLLKNIDERGKGTVSR |

| | | |
|---|---|---|
| | GCTGAAAAATATCGACGAGCGCGGAAAAGGCACCGTCAGCCGTGACTGGGT<br>TGGTCGCGACGCAGGCAAAATCGCGGCGGCAATCGGCCTTAAAGTTCCGCA<br>AGAAACGCGCCTGCTGTTTGTGGAAACCACCGCAGAACATCCGTTTGCCGTG<br>ACTGAACTGATGATGCCGGTGTTGCCCGTCGTGCGCGTCGCCAACGTGGCGG<br>ATGCCATTGCGCTAGCGGTGAAACTGGAAGGCGGTTGCCACCACACGGCGG<br>CAATGCACTCGCGCAACATCGAAAACATGAACCAGATGGCGAATGCTATTG<br>ATACCAGCATTTTCGTTAAGAACGGACCGTGCATTGCCGGGCTGGGGCTGGG<br>CGGGGAAGGCTGGACCACCATGACCATCACCACGCCAACCGGTGAAGGGGT<br>AACCAGCGCGCGTACGTTTGTCCGTCTGCGTCGCTGTGTATTAGTCGATGCG<br>TTTCGCATTGTTTAA | DWVGRDAGKIAAAIGLK<br>VPQETRLLFVETTAEHPFA<br>VTELMMPVLPVVRVANV<br>ADAIALAVKLEGGCHHTA<br>AMHSRNIENMNQMANAI<br>DTSIFVKNGPCIAGLGLGG<br>EGWTTMTITTPTGEGVTS<br>ARTFVRLRRCVLVDAFRI<br>V* |
| pACYC-eutG | ATGCAAAATGAATTGCAGACCGCGCTCTTTCAGGCGTTCGATACCCTGAATC<br>TGCAACGGGTAAAAACATTTAGCGTTCCACCGGTGACGCTTTGCGGTCCGGG<br>CTCGGTGAGCAGTTGCGGACAGCAAGCGCAAACGCGTGGGCTGAAACATCT<br>GTTCGTGATGGCAGACAGCTTTTTGCATCAGGCAGGGATGACCGCCGGGCTG<br>ACGCGTAGCCTGACCGTTAAAGGTATCGCCATGACGCTCTGGCCATGTCCGG<br>TGGGCGAACCGTGCATTACCGACGTGTGTGCAGCCGTGGCGCAGTTGCGTG<br>AGTCAGGCTGTGATGGGGTGATCGCGTTTGGCGGCGGCTCGGTGCTGGATGC<br>GGCGAAAGCCGTGACGTTGCTGGTGACGAACCCGGATAGCACGCTGGCAGA<br>GATGTCAGAAACCAGCGTTCTGCAACCGCGCTTGCCGCTGATTGCCATTCCA<br>ACTACCGCCGGAACCGGCTCTGAAACCACCAATGTAACGGTGATTATCGAC<br>GCGGTGAGCGGGCGCAAGCAGGTGTTAGCCCATGCCTCGCTGATGCCGGAT<br>GTGGCGATCCTCGACGCCGCATTGACCGAAGGTGTGCCGTCGCATGTCACGG<br>CGATGACCGGCATTGATGCGTTAACCCATGCCATTGAAGCATACAGCGCCCT<br>GAACGCTACACCGTTTACCGACAGTCTGGCGATTGGTGCCATTGCGATGATT<br>GGCAAATCGCTGCCGAAAGCGGTGGGCTACGGTCACGACCTTGCCGCGCGC<br>GAGAGCATGTTGCTGGCTTCATGTATGGCGGGAATGGCGTTTTCCAGTGCGG<br>GTCTTGGGTTGTGCCACGCGATGGCGCATCAGCCGGGCGCGGCGCTGCATAT<br>TCCGCACGGTCTCGCGAACGCCATGTTGCTGCCAACGGTGATGGAATTTAAC<br>CGGATGGTTTGTCGTGAACGCTTTAGTCAGATTGGTCGGGCACTGCGAACTA<br>AAAAATCCGACGATCGTGACGCTATTAACGCGGTAAGTGAGCTGATTGCGG<br>AAGTTGGGATTGGTAAACGACTGGGCGATGTTGGTGCGACATCTGCGCATTA<br>CGGCGCATGGGCGCAGGCCGCGCTGGAAGATATTTGTCTGCGCAGTAACCC<br>GCGTACCGCCAGCCTGGAGCAGATTGTCGGCCTGTACGCAGCGGCGCAATA<br>A | MQNELQTALFQAFDTLNL<br>QRVKTFSVPPVTLCGPGS<br>VSSCGQQAQTRGLKHLFV<br>MADSFLHQAGMTAGLTR<br>SLTVKGIAMTLWPCPVGE<br>PCITDVCAAVAQLRESGC<br>DGVIAFGGGSVLDAAKAV<br>TLLVTNPDSTLAEMSETS<br>VLQPRLPLIAIPTTAGTGSE<br>TTNVTVIIDAVSGRKQVL<br>AHASLMPDVAILDAALTE<br>GVPSHVTAMTGIDALTHA<br>IEAYSALNATPFTDSLAIG<br>AIAMIGKSLPKAVGYGHD<br>LAARESMLLASCMAGMA<br>FSSAGLGLCHAMAHQPG<br>AALHIPHGLANAMLLPTV<br>MEFNRMVCRERFSQIGRA<br>LRTKKSDDRDAINAVSELI<br>AEVGIGKRLGDVGATSAH<br>YGAWAQAALEDICLRSNP<br>RTASLEQIVGLYAAAQ* |
| pACYC-fucO | ATGGCTAACAGAATGATTCTGAACGAAACGGCATGGTTTGGTCGGGGTGCT<br>GTTGGGGCTTTAACCGATGAGGTGAAACGCCGTGGTTATCAGAAGGCGCTG<br>ATCGTCACCGATAAAACGCTGGTGCAATGCGGCGTGGTGGCGAAAGTGACC<br>GATAAGATGGATGCTGCAGGGCTGGCATGGGCGATTTACGACGGCGTAGTG<br>CCCAACCCAACAATTACTGTCGTCAAAGAAGGGCTCGGTGTATTCCAGAATA<br>GCGGCGCGGATTACCTGATCGCTATTGGTGGTGGTTCTCCACAGGATACTTG<br>TAAAGCGATTGGCATTATCAGCAACAACCCGGAGTTTGCCGATGTGCGTAGC<br>CTGGAAGGGCTTTCCCCGACCAATAAACCCAGTGTACCGATTCTGGCAATTC<br>CTACCACAGCAGGTACTGCGGCAGAAGTGACCATTAACTACGTGATCACTG<br>ACGAAGAGAAACGGCGCAAGTTTGTTTGCGTTGATCCGCATGATATCCCGCA<br>GGTGGCGTTTATTGACGCTGACATGATGGATGGTATGCCTCCAGCGCTGAAA<br>GCTGCGACGGGTGTCGATGCGCTCACTCATGCTATTGAGGGGTATATTACCC<br>GTGGCGCGTGGGCGCTAACCGATGCACTGCACATTAAAGCGATTGAAATCA<br>TTGCTGGGGCGCTGCGAGGATCGGTTGCTGGTGATAAGGATGCCGGAGAAG<br>AAATGGCGCTCGGGCAGTATGTTGCGGGTATGGGCTTCTCGAATGTTGGGTT<br>AGGGTTGGTGCATGGTATGGCGCATCCACTGGGCGCGTTTTATAACACTCCA<br>CACGGTGTTGCGAACGCCATCCTGTTACCGCATGTCATGCGTTATAACGCTG<br>ACTTTACCGGTGAGAAGTACCGCGATATCGCGCGCGTTATGGGCGTGAAAG<br>TGGAAGGTATGAGCCTGGAAGAGGCGCGTAATGCCGCTGTTGAAGCGGTGT<br>TTGCTCTCAACCGTGATGTCGGTATTCCGCCACATTTGCGTGATGTTGGTGTA<br>CGCAAGGAAGACATTCCGGCACTGGCGCAGGCGGCACTGGATGATGTTTGT<br>ACCGGTGGCAACCCGCGTGAAGCAACGCTTGAGGATATTGTAGAGCTTTAC<br>CATACCGCCTGGTAA | MANRMILNETAWFGRGA<br>VGALTDEVKRRGYQKALI<br>VTDKTLVQCGVVAKVTD<br>KMDAAGLAWAIYDGVVP<br>NPTITVVKEGLGVFQNSG<br>ADYLIAIGGGSPQDTCKAI<br>GIISNNPEFADVRSLEGLSP<br>TNKPSVPILAIPTTAGTAA<br>EVTINYVITDEEKRRKFVC<br>VDPHDIPQVAFIDADMMD<br>GMPPALKAATGVDALTH<br>AIEGYITRGAWALTDALHI<br>KAIEIIAGALRGSVAGDKD<br>AGEEMALGQYVAGMGFS<br>NVGLGLVHGMAHPLGAF<br>YNTPHGVANAILLPHVMR<br>YNADFTGEKYRDIARVM<br>GVKVEGMSLEEARNAAV<br>EAVFALNRDVGIPPHLRD<br>VGVRKEDIPALAQAALDD<br>VCTGGNPREATLEDIVEL<br>YHTAW* |
| pACYC-gldA | ATGGACCGCATTATTCAATCACCGGGTAAATACATCCAGGGCGCTGATGTGA<br>TTAATCGTCTGGGCGAATACCTGAAGCCGCTGGCAGAACGCTGGTTAGTGGT<br>GGGTGACAAATTTGTTTTAGGTTTTGCTCAATCCACTGTCGAGAAAAGCTTT<br>AAAGATGCTGGACTGGTAGTAGAAATTGCGCCGTTTGGCGGTGAATGTTCGC<br>AAAATGAGATCGACCGTCTGCGTGGCATCGCGGAGACTGCGCAGTGTGGCG<br>CAATTCTCGGTATCGGTGGCGGAAAAACCCTCGATACTGCCAAAGCACTGG<br>CACATTTCATGGGTGTTCCGGTAGCGATCGCACCGACTATCGCCTCTACCGA<br>TGCACCGTGCAGCGCATTGTCTGTTATCTACACCGATGAGGGTGAGTTTGAC<br>CGCTATCTGCTGTTGCCAAATAACCCGAATATGGTCATTGTCGACACCAAAA<br>TCGTCGCTGGCGCACCTGCACGTCTGTTAGCGGCGGGTATCGGCGATGCGCT<br>GGCAACCTGGTTTGAAGCGCGTGCCTGCTCTCGTAGCGGCGCGACCACCATG<br>GCGGGCGGCAAGTGCACCCAGGCTGCGCTGGCACTGGCTGAACTGTGCTAC<br>AACACCCTGCTGGAAGAAGGCGAAAAAGCGATGCTTGCTGCCGAACAGCAT | MDRIIQSPGKYIQGADVIN<br>RLGEYLKPLAERWLVVG<br>DKFVLGFAQSTVEKSFKD<br>AGLVVEIAPFGGECSQNEI<br>DRLRGIAETAQCGAILGIG<br>GGKTLDTAKALAHFMGV<br>PVAIAPTIASTDAPCSALS<br>VIYTDEGEFDRYLLLPNNP<br>NMVIVDTKIVAGAPARLL<br>AAGIGDALATWFEARACS<br>RSGATTMAGGKCTQAAL<br>ALAELCYNTLLEEGEKAM<br>LAAEQHVVTPALERVIEA |

| | | |
|---|---|---|
| | GTAGTGACTCCGGCGCTGGAGCGCGTGATTGAAGCGAACACCTATTTGAGC<br>GGTGTTGGTTTTGAAAGTGGTGGTCTGGCTGCGGCGCACGCAGTGCATAACG<br>GCCTGACCGCTATCCCGGACGCGCATCACTATTATCACGGTGAAAAAGTGGC<br>ATTCGGTACGCTGACGCAGCTGGTTCTGGAAAATGCGCCGGTGGAGGAAAT<br>CGAAACCGTAGCTGCCCTTAGCCATGCGGTAGGTTTGCCAATAACTCTCGCT<br>CAACTGGATATTAAAGAAGATGTCCCGGCGAAAATGCGAATTGTGGCAGAA<br>GCGGCATGTGCAGAAGGTGAAACCATTCACAACATGCCTGGCGGCGCGACG<br>CCAGATCAGGTTTACGCCGCTCTGCTGGTAGCCGACCAGTACGGTCAGCGTT<br>TCCTGCAAGAGTGGGAATAA | NTYLSGVGFESGGLAAAH<br>AVHNGLTAIPDAHHYYH<br>GEKVAFGTLTQLVLENAP<br>VEEIETVAALSHAVGLPIT<br>LAQLDIKEDVPAKMRIVA<br>EAACAEGETIHNMPGGAT<br>PDQVYAALLVADQYGQR<br>FLQEWE* |
| pACYC-gpr | ATGGTCTGGTTAGCGAATCCCGAACGTTACGGGCAGATGCAATACCGCTATT<br>GCGGAAAAAGTGGTTTACGCCTGCCCGCGTTATCGCTCGGTTTATGGCACAA<br>TTTCGGTCACGTTAACGCGCTGGAATCACAGCGTGCGATCCTGCGTAAAGCG<br>TTTGATTTGGGCATTACGCACTTTGATTTAGCCAACAATTACGGGCCGCCTC<br>CAGGAAGCGCAGAAGAGAACTTTGGTCGCCTGCTGCGGGAGGATTTTGCCG<br>CTTATCGCGATGAACTGATTATCTCTACCAAGGCTGGCTACGATATGTGGCC<br>CGGCCCTTACGGCTCTGGCGGTTCACGTAAATACCTGCTCGCCAGCCTCGAC<br>CAAAGCCTGAAGCGTATGGGGCTTGAGTATGTCGATATCTTTTACTCTCATC<br>GCGTCGATGAAAATACGCCGATGGAAGAAACCGCCTCTGCGCTGGCTCATG<br>CGGTACAAAGCGGTAAGGCGCTGTATGTCGGGATCTCCTCTTACTCGCCAGA<br>GCGGACGCAAAAAATGGTCGAGTTGCTGCGCGAGTGGAAAATTCCGCTGTT<br>AATTCATCAACCTTCGTACAATTTACTGAACCGCTGGGTGGATAAAAGCGGC<br>CTGCTGGATACCCTGCAAAATAACGGCGTGGGCTGTATTGCCTTTACTCCTC<br>TGGCTCAGGGATTGCTGACCGGAAAATATCTCAACGGCATTCCGCAAGATTC<br>ACGGATGCATCGTGAAGGGAATAAAGTTCGTGGTCTGACACCGAAAATGCT<br>TACCGAAGCCAACCTCAACAGCCTGCGCTTATTGAATGAAATGGCACAGCA<br>GCGTGGACAATCAATGGCGCAAATGGCGTTAAGCTGGTTGCTGAAAGATGA<br>TCGCGTGACGTCGGTATTGATTGGTGCCAGCCGCGCGGAGCAACTAGAGGA<br>GAACGTGCAGGCGCTGAATAATCTGACATTTAGCACCAAGGAGCTGGCGCA<br>GATTGATCAGCATATCGCCGATGGCGAGCTGAATCTGTGGCAGGCGTCTTCC<br>GATAAATGA | MVWLANPERYGQMQYR<br>YCGKSGLRLPALSLGLWH<br>NFGHVNALESQRAILRKA<br>FDLGITHFDLANNYGPPPG<br>SAEENFGRLLREDFAAYR<br>DELIISTKAGYDMWPGPY<br>GSGGSRKYLLASLDQSLK<br>RMGLEYVDIFYSHRVDEN<br>TPMEETASALAHAVQSGK<br>ALYVGISSYSPERTQKMV<br>ELLREWKIPLLIHQPSYNL<br>LNRWVDKSGLLDTLQNN<br>GVGCIAFTPLAQGLLTGK<br>YLNGIPQDSRMHREGNKV<br>RGLTPKMLTEANLNSLRL<br>LNEMAQQRGQSMAQMA<br>LSWLLKDDRVTSVLIGAS<br>RAEQLEENVQALNNLTFS<br>TKELAQIDQHIADGELNL<br>WQASSDK* |
| pACYC-ybbO | ATGACTCATAAAGCAACGGAGATCCTGACAGGTAAAGTTATGCAAAAATCG<br>GTCTTAATTACCGGATGTTCCAGTGGAATTGGCCTGGAAAGCGCGCTCGAAT<br>TAAAACGCCAGGGTTTTCATGTGCTGGCAGGTTGCCGGAAACCGGATGATGT<br>TGAGCGCATGAACAGCATGGGATTTACCGGCGTGTTGATCGATCTGGATTCA<br>CCAGAAAGTGTTGATCGCGCAGCCGACGAGGTGATCGCCCTGACCGATAAT<br>TGTCTGTATGGGATCTTTAACAATGCCGGATTCGGCATGTATGGCCCCCTTTC<br>CACCATCAGCCGTGCGCAGATGGAACAGCAGTTTTCCGCCAACTTTTTCGGC<br>GCACACCAGCTCACCATGCGCCTGTTACCCGCGATGTTACCGCACGGTGAAG<br>GGCGTATTGTGATGACATCATCGGTGATGGGATTAATCTCCACGCCGGGTCG<br>TGGCGCTTACGCGGCCAGTAAATATGCGCTGGAGGCGTGGTCAGATGCACT<br>GCGCATGGAGCTGCGCCACAGCGGAATTAAAGTCAGCCTGATCGAACCCGG<br>TCCCATTCGTACTCGCTTCACCGACAACGTCAACCAGACGCAAAGTGATAAA<br>CCAGTCGAAAATCCCGGCATCGCCGCCCGCTTTACGTTGGGACCGGAAGCG<br>GTGGTGGACAAAGTACGCCATGCTTTTATTAGCGAGAAGCCGAAGATGCGC<br>TATCCGGTGACGCTGGTGACCTGGGCGGTAATGGTGCTTAAGCGCCTGCTGC<br>CGGGGCGCGTGATGGACAAAATATTGCAGGGGTGA | MTHKATEILTGKVMQKS<br>VLITGCSSGIGLESALELK<br>RQGFHVLAGCRKPDDVE<br>RMNSMGFTGVLIDLDSPE<br>SVDRAADEVIALTDNCLY<br>GIFNNAGFGMYGPLSTISR<br>AQMEQQFSANFFGAHQLT<br>MRLLPAMLPHGEGRIVMT<br>SSVMGLISTPGRGAYAAS<br>KYALEAWSDALRMELRH<br>SGIKVSLIEPGPIRTRFTDN<br>VNQTQSDKPVENPGIAAR<br>FTLGPEAVVDKVRHAFISE<br>KPKMRYPVTLVTWAVMV<br>LKRLLPGRVMDKILQG* |
| -pACYC-yghA | ATGTCTCATTTAAAAGACCCGACCACGCAGTATTACACTGGTGAATATCCCA<br>AACAGAAACAACCGACGCCAGGCATCCAGGCGAAGATGACACCGGTACCG<br>GATTGCGGCGAGAAAACCTATGTTGGTAGCGGTCGCCTGAAAGATCGTAAA<br>GCACTGGTGACAGGGGGCGATTCCGGAATAGGTCGCGCTGCCGCCATCGCT<br>TACGCGCGTGAAGGGGCTGACGTGGCGATCAGTTATCTTCCCGTGGAAGAA<br>GAAGACGCTCAGGATGTGAAAAAGATCATTGAAGAATGCGGACGCAAAGCC<br>GTTCTGCTGCCAGGCGATTTAAGCGATGAGAAATTTGCCCGTTCGCTGGTTC<br>ACGAAGCGCACAAGGCGTTAGGCGGGCTGGATATTATGGCGCTGGTCGCCG<br>GGAAACAGGTTGCCATTCCGGATATTGCAGACCTCACCAGCGAACAGTTTCA<br>AAAGACCTTTGCCATTAACGTTTTCGCGCTGTTCTGGCTAACCCAGGAAGCG<br>ATCCCCCTGCTACCGAAAGGTGCAAGTATCATCACCACTTCGTCAATCCAGG<br>CATACCAGCCAAGTCCGCATTTACTGGACTATGCGGCTACGAAGGCGGCGA<br>TTCTGAACTACAGCCGTGGCTTGGCAAAACAGGTCGCGGAGAAAGGTATTC<br>GGGTGAATATTGTCGCGCCAGGCCCGATCTGGACAGCACTGCAAATTTCCGG<br>CGGACAAACGCAGGATAAGATCCCGCAGTTTGGTCAGCAAACGCCGATGAA<br>ACGTGCGGGGCAACCGGCGGAACTGGCCCCTGTATATGTTTATCTGGCAAGT<br>CAGGAGTCGAGCTACGTCACCGCAGAAGTGCACGGCGTGTGCGGCGGCGAG<br>CATTTAGGTTAA | MSHLKDPTTQYYTGEYPK<br>QKQPTPGIQAKMTPVPDC<br>GEKTYVGSGRLKDRKAL<br>VTGGDSGIGRAAAIAYAR<br>EGADVAISYLPVEEEDAQ<br>DVKKIIEECGRKAVLLPG<br>DLSDEKFARSLVHEAHKA<br>LGGLDIMALVAGKQVAIP<br>DIADLTSEQFQKTFAINVF<br>ALFWLTQEAIPLLPKGASII<br>TTSSIQAYQPSPHLLDYAA<br>TKAAILNYSRGLAKQVAE<br>KGIRVNIVAPGPIWTALQI<br>SGGQTQDKIPQFGQQTPM<br>KRAGQPAELAPVYVYLAS<br>QESSYVTAEVHGVCGGEH<br>LG* |
| pACYC-yiaY | ATGGCAGCTTCAACGTTCTTTATTCCTTCTGTGAATGTCATCGGCGCTGATTC<br>ATTGACTGATGCAATGAATATGATGGCAGATTATGGATTTACCCGTACCTTA<br>ATTGTCACTGACAATATGTTAACGAAATTAGGTATGGCGGGCGATGTGCAA<br>AAAGCACTGGAAGAACGCAATATTTTTAGCGTTATTTATGATGGCACCCAAC<br>CTAACCCCACCACGGAAAACGTCGCCGCAGGTTTGAAATTACTTAAAGAGA<br>ATAATTGCGATAGCGTGATCTCCTTAGGCGGTGGTTCTCCACACGACTGCGC | MAASTFFIPSVNVIGADSL<br>TDAMNMMADYGFTRTLI<br>VTDNMLTKLGMAGDVQK<br>ALEERNIFSVIYDGTQPNP<br>TTENVAAGLKLLKENNCD<br>SVISLGGGSPHDCAKGIAL |

| | | |
|---|---|---|
| | AAAAGGTATTGCGCTGGTGGCAGCCAATGGCGGCGATATTCGCGATTACGA<br>AGGCGTTGACCGCTCTGCAAAACCGCAGCTGCCGATGATCGCCATCAATACC<br>ACGGCGGGTACGGCCTCTGAAATGACCCGTTTCTGCATCATCACTGACGAAG<br>CGCGTCATATCAAAATGGCGATTGTTGATAAACATGTCACTCCGCTGCTTTC<br>TGTCAATGACTCCTCTCTGATGATTGGTATGCCGAAGTCACTGACCGCCGCA<br>ACGGGTATGGATGCCTTAACGCACGCTATCGAAGCATATGTTTCTATTGCCG<br>CCACGCCGATCACTGACGCTTGTGCACTGAAAGCCGTGACCATGATTGCCGA<br>AAACCTGCCGTTAGCCGTTGAAGATGGCAGTAATGCGAAAGCGCGTGAAGC<br>AATGGCTTATGCCCAGTTCCTCGCCGGTATGGCGTTCAATAATGCTTCTCTG<br>GGTTATGTTCATGCGATGGCGCACCAGCTGGGCGGTTTCTACAACCTGCCAC<br>ACGGTGTATGTAACGCCGTTTTGCTGCCGCACGTTCAGGTATTCAACAGCAA<br>AGTCGCCGCTGCACGTCTGCGTGACTGTGCCGCTGCAATGGGCGTGAACGTG<br>ACAGGTAAAAACGACGCGGAAGGTGCTGAAGCCTGCATTAACGCCATCCGT<br>GAACTGGCGAAGAAAGTGGATATCCCGGCAGGCCTACGCGACCTGAACGTG<br>AAAGAAGAAGATTTCGCGGTATTGGCGACTAATGCCCTGAAAGATGCCTGT<br>GGCTTTACTAACCCGATCCAGGCAACTCACGAAGAAATTGTGGCGATTTATC<br>GCGCAGCGATGTAA | VAANGGDIRDYEGVDRS<br>AKPQLPMIAINTTAGTASE<br>MTRFCIITDEARHIKMAIV<br>DKHVTPLLSVNDSSLMIG<br>MPKSLTAATGMDALTHAI<br>EAYVSIAATPITDACALKA<br>VTMIAENLPLAVEDGSNA<br>KAREAMAYAQFLAGMAF<br>NNASLGYVHAMAHQLGG<br>FYNLPHGVCNAVLLPHVQ<br>VFNSKVAAARLRDCAAA<br>MGVNVTGKNDAEGAEAC<br>INAIRELAKKVDIPAGLRD<br>LNVKEEDFAVLATNALKD<br>ACGFTNPIQATHEEIVAIY<br>RAAM* |

**Table S4.** Genes targeted for translational knockout and the position of premature stop codon mutagenesis.

| Gene | Enzyme family | Total translated length (amino acids) | Active site residue | Positions of inserted premature stop codon (targeted codon in parenthesis) | Length of truncated protein (amino acids) |
|---|---|---|---|---|---|
| *adhP* | ADH | 336 | | 87 (TAA), 88 (TGA) | 86 |
| *fucO* | ADH | 382 | 267 [3,4] | 29 (TGA), 30 (TAA) | 28 |
| *eutG* | ADH | 395 | | 41 (TGA), 42 (TAA),43 (TAA) | 40 |
| *yiaY* | ADH | 383 | | 82 (TAA), 83 (TGA) | 81 |
| *adhE* | ADH | 891 | 246 (homology prediction) | 21 (TGA), 22 (TAA) | 20 |
| *eutE* | ADH | 467 | | 45 (TAG), 48 (TAA), 49 (TGA) | 44 |
| *gldA* | ADH | 367 | 245 [5] | 67 (TAA), 68 (TGA), 69 (TAG) | 66 |
| *gpr* | AKR | 346 | 138 [6] | 43 (TAA), 44 (TGA), 45 (TAG) | 42 |
| *ybbO* | ADH | 269 | 146 (homology prediction) | 15 (TAA), 16 (TGA), 17 (TAG) | 14 |
| *yghA* | ADH | 294 | 199 (homology prediction) | 3 (TAG), 4 (TGA), 5 (TAA) | 2 |

### II. Supplementary Methods

#### Strains and plasmids

*Escherichia coli* strains and plasmids used are listed in **Table S1**. Molecular cloning and vector propagation were performed in DH5α (NEB). Polymerase chain reaction (PCR) based DNA replication was performed using KOD XTREME Hot Start Polymerase (MilliporeSigma) for plasmid backbones. Cloning was performed using Gibson Assembly. Oligos for PCR amplification and translational knockouts are shown in **Table S2**. Oligos were purchased from Integrated DNA Technologies (IDT). The DNA sequence and translated sequence of proteins overexpressed in this paper are found in **Table S3**. The pORTMAGE-Ec1 recombineering plasmid was kindly provided by Timothy Wannier and George Church of Harvard Medical School.

**Chemicals**

The following compounds were purchased from MilliporeSigma: sodium borate decahydrate, sodium phosphate dibasic anhydrous, chloramphenicol, kanamycin sulfate, dimethyl sulfoxide (DMSO), boric acid, L-alanine, and HEPES. D-glucose and m-toluic acid. The following compounds were purchased from Alfa Aesar: agarose and ethanol were purchased. The following compounds were purchased from Fisher Scientific: isopropyl ß-D-1-thiogalactopyranoside (IPTG), acetonitrile, sodium chloride, trifluoroacetic acid, LB Broth powder (Lennox), and LB Agar powder (Lennox). A MOPS EZ rich defined medium kit was purchased from Teknova. Taq DNA ligase was purchased from GoldBio. Anhydrotetracycline (aTc) was purchased from Cayman Chemical. Phusion DNA polymerase and T5 exonuclease were purchased from New England BioLabs (NEB). Sybr Safe DNA gel stain was purchased from Invitrogen. The following compounds were purchased from TCI America: Pyridoxal 5'-phosphate (PLP), *ortho*-phthalaldehyde and 3-mercaptopropionic acid

**Cloning of TA and AlaDH**

Molecular cloning and vector propagation were performed in *E. coli* DH5α. Codon-optimized genes were PCR amplified using KOD Xtreme polymerase The ω-transaminase from *Chromobacterium violaceum* (CvTA, WP_011135573.1) and alanine dehydrogenase from *Bacillus subtilis* (BsAlaDH, WP_003243280.1 ) were purchased as gene fragments from IDT and cloned using Gibson Assembly into a pACYC vector harboring an N-terminal $His_6$-tag for purification, p15a ori, and chloramphenicol acetyltransferase gene for antibiotic selection. All plasmids were verified by Sanger sequencing.

**Cloning of RARE.Δ16 gene KO**

The targeted genes for inactivation in RARE.Δ16 (*adhP*, *fucO*, *eutG*, *yiaY*, *adhE*, *eutE*, *gldA*, *gpr*, *ybbO* and *yghA*) were amplified by PCR from the *E. coli* MG1655 genome and were cloned into a pACYC p15a ori, and chloramphenicol acetyltransferase gene for antibiotic selection by Gibson Assembly.

**Culture Conditions**

Cultures were grown in LB-Lennox medium (LB: 10 g/L bacto tryptone, 5 g/L sodium chloride, 5 g/L yeast extract), M9-glucose minimal media[7] with Corning® Trace Elements A (1.60 µg/mL $CuSO_4 \cdot 5H_2O$, 863.00 µg/mL $ZnSO_4 \cdot 7H_2O$, 17.30 µg/mL Selenite • 2Na, 1155.10 µg/mL ferric citrate) and 1.5% glucose, or MOPS EZ rich defined media (Teknova M2105) with 2% glucose (MOPS EZ Rich-glucose).

**Stability Assays**

For testing stability in metabolically active cells, cultures of each *E. coli* strain to be tested were inoculated from a frozen stock and grown to confluence overnight in 5 mL of LB media. Confluent overnight cultures were then used to inoculate experimental cultures to an initial starting $OD_{600}$ of 0.01 in 300 µL volumes in a 96-deep-well plate (Thermo Scientific™ 260251). Cultures were supplemented with 5 mM of TPAL (prepared in 100 mM stocks in 100% DMSO) at mid-exponential phase ($OD_{600}$ = 0.5-0.8). Cultures were incubated at 37 °C with shaking at 1000 RPM and an orbital radius of 3 mm. Samples were taken by pipetting 200 µL from the cultures, centrifuging in a different 96-deep-well plate and collecting the extracellular broth. Compounds were quantified over a 24 h period using HPLC with samples collected at 4 h and 24 h.

For metabolically active cell stability testing with targeted single gene knockouts plasmids, cultures of *E. coli* RARE.Δ6 and RARE.Δ16 harboring a gene from the 10 targeted knockouts were inoculated from a frozen stock and grown to confluence overnight in 5 mL of LB media with 34 µg/mL chloramphenicol. Confluent overnight cultures were then used to inoculate experimental cultures with cm to an initial starting $OD_{600}$ of 0.01 in 600 µL volumes in a 96-deep-well plate. Cultures were supplemented with 5 mM of heterologous aldehydes at mid-exponential phase ($OD_{600}$ = 0.5-0.8). Cultures were incubated at 37 °C with shaking at 1000 RPM and an orbital radius of 3 mm. Samples were taken by pipetting 100 µL from the cultures, centrifuging in a different 96-deep-well plate and collecting the extracellular broth. Compounds were quantified over a 24 h period using HPLC with samples collected at 4 h and 24 h.

**RNAseq**

Total RNA was sampled from duplicate cultures for each condition. To prepare samples for RNA extraction, cultures of *E. coli* RARE.Δ6 were inoculated at a 1:50 dilution from a confluent overnight culture into 3 mL of M9-glucose minimal media in 14 mL culture tubes. Cultures were then grown at 37 °C for 2 h (until $OD_{600}$~0.4 was reached). After 2 h, 1 mM of TPAL was supplemented to cultures from 400 mM stocks prepared in DMSO. Culture tubes were then snapped shut and sealed with parafilm to limit loss of aldehyde due to volatility. Cultures were then grown at 37 °C for an additional 1.5 h. Then, 5 x $10^8$ cells (as measured by optical density) were mixed with RNAprotect Bacteria Reagent (1:2 volume ratio). Then, bacterial lysis and RNA purification were performed using a Qiagen RNAeasy purification kit (Handbook 2020). Following purification to ensure better purity, the RNA was precipitated by adding 6 µL if 5 M sodium acetate with 180 µL of ethanol. Then, the samples were placed at -20 °C. After 18 h, the samples were centrifuged at 18,000 x g at 4 °C for 10 min and then the supernatant was decanted. After the samples dried, they were resuspended in RNAse free water and frozen at -80 °C until submission to Novogene for total RNA sequencing. Transcriptional fold change was performed with TPAL supplementation compared to a RARE.Δ6 baseline in duplicates.

**Translational genomic knockouts**

Translational knockouts to the strain were performed using 10 rounds of multiplexed automatable genome engineering (MAGE) where stop codons were introduced into the genomic sequence for each aldehyde dehydrogenase target in the upstream portion of the gene (Table.S4). Construction of RARE.Δ16strain was done with two subsets of knockouts (subset #1:*adhP*, *fucO*, *eutG*, *yiaY* and *adhE*, subset #2: *eutE*, *gldA*, *gpr*, *ybbO*, and *yghA*) performed in the RARE.Δ6 strain[1]. Construction of RARE.Δ10 was done using the same oligos that were used for the RARE.Δ16 strain. MAGE was performed using the pORTMAGE-Ec1 recombineering plasmid[2]. Briefly, bacterial cultures were inoculated with 1:100 dilution in 3 mL of LB media with 30 μg/mL kanamycin (Kan) and grown at 37 °C until $OD_{600}$ of 0.4 - 0.6 was reached. Then, the proteins responsible for recombineering were induced with 1 mM m-toluic acid and cultures were grown at 37 °C for an additional 15 min. Cells were then prepared for electroporation by washing 1 mL three times with refrigerated 10% glycerol and then resuspending in 50 µL of 10% glycerol with each knockout oligo within a subset added at 1 µM. Cells were then electroporated and recovered in 3 mL of LB with 30 µg/mL kanamycin to be used for subsequent rounds. Preliminary assessment of knockouts was performed using mascPCR (multiplexed allele specific colony PCR)[8] and confirmed using Sanger Sequencing. RARE.Δ11, RARE.Δ16 and RARE.Δ10 were cured of the pORTMAGE-Ec1 plasmid following confirmation of genomic knockouts.

**Growth rate and protein production assay**

For growth rate testing, confluent overnight cultures were used to inoculate experimental cultures in LB and MOPS EZ Rich media at 100x dilution in 200 µL volumes in a Greiner black clear bottom 96 well plate (Greiner 655090). Cultures were grown for 12 h in a Spectramax i3x plate reader with medium plate shaking at 37 °C with absorbance readings at 600 nm taken every 7 min to determine growth rate of each strain. For protein expression testing, strains were transformed with a pZE-Ub-sfGFP reporter plasmid. Confluent cultures of cells harboring the pZE-Ub-sfGFP were grown as described above with 30 μg/mL kanamycin and 0.2 nM aTc added at inoculation.

***In vivo* transaminase activity**

For metabolically active ω-transaminase testing, cultures of MG1655, RARE.Δ6 and RARE.Δ16 harboring a plasmid containing both CvTA and AlaDH were inoculated from a frozen stock and grown to confluence overnight in 5 mL of LB media with 34 μg/mL chloramphenicol. Confluent overnight cultures were then used to inoculate experimental cultures in a 96-deep-well plate initial starting $OD_{600}$ of 0.01 in 300 μL in LB with34 μg/mL chloramphenicol. Cultures were supplemented with 1 mM of IPTG, 100 mM L-alanine (amine donor) and with 5 mM of TPAL (prepared in 100 mM stocks in DMSO) at mid-exponential phase ($OD_{600}$ = 0.5-0.8). Cultures were incubated at 37 °C with shaking at 1000 RPM and an orbital radius of 3 mm for 4 hours. Samples were taken by centrifuging in a 96-deep-well plate and collecting the extracellular broth. Compounds were quantified after 24 h using HPLC.

**HPLC analysis**

TPAL, 4HMB, 4HMBA and pXYL were quantified using reverse-phase high-performance liquid chromatography (HPLC) with an Agilent 1260 Infinity with a Zorbax Eclipse Plus-C18 column with a guard column installed. Amines were derivatized with *ortho*-pthalaldehyde and 3-mercaptopropionic acid and identified using reverse-phase high-performance liquid chromatography (RP-HPLC) with an Agilent 1260 Infinity with a Zorbax Eclipse Plus-C18 column with a guard column installed. The methods for detection of TPAL, 4HMB, 4HMBA and pXYL used were described in Gopal et al[9].
